# Insights into the bioluminescence systems of three sea pens (Cnidaria: Anthozoa): from *de novo* transcriptome analyses to biochemical assays

**DOI:** 10.1101/2024.04.30.591678

**Authors:** Laurent Duchatelet, Gabriela A. Galeazzo, Constance Coubris, Laure Bridoux, René Rezsohazy, Marcelo R.S. Melo, Martin Marek, Danilo T. Amaral, Sam Dupont, Anderson G. Oliveira, Jérôme Delroisse

## Abstract

Bioluminescence is the production of visible light by living organisms. It occurs through the oxidation of a luciferin substrate catalysed by luciferase enzymes. Auxiliary proteins, such as fluorescent proteins and luciferin-binding proteins, can modify the light emitted wavelength or stabilize reactive luciferin molecules, respectively. Additionally, calcium ions are crucial for the luminescence across various species. Despite the large phylogenetic distribution of bioluminescent organisms, only a few systems have been comprehensively studied. Notably, cnidarian species of the *Renilla* genus utilize a coelenterazine-dependent luciferase, a calcium-dependent coelenterazine-binding protein and a green fluorescent protein. We investigated the bioluminescence of three sea pen species: *Pennatula phosphorea, Anthoptilum murrayi* and *Funiculina quadrangularis* (Pennatuloidea, Anthozoa). Their light-emission spectra reveal peaks at 510, 513 and 485 nm, respectively. A coelenterazine-based reaction was demonstrated in all three species. Using transcriptome analyses, we identified predicted transcripts coding for luciferases, green fluorescent proteins and coelenterazine-binding proteins for *P. phosphorea* and *A. murrayi*. Immunodetection confirmed the expression of luciferase in *P. phosphorea* and *F. quadrangularis*. We also expressed recombinant luciferase of *A. murrayi*, confirming its activity. We highlighted the role of calcium ions in bioluminescence, possibly associated with the mechanism of substrate release at the level of coelenterazine-binding proteins. The study proposes a model for anthozoan bioluminescence, offering new avenues for future ecological and functional research on these luminous organisms.

## Introduction

Bioluminescence, the production of visible light by a living organism, involves the oxidation of a luciferin substrate catalyzed by a luciferase enzyme. In some bioluminescent systems, luciferase and luciferin form a stable complex known as “photoprotein”, requiring additional co-factors to be functional [1]. Luciferases *sensu lato* (*i.e.,* luciferases *stricto sensu* and photoproteins) are classically considered taxon-specific (*i.e.,* each clade is characterized by its own luciferase) [2]. Nevertheless, homologous luciferases can sometimes be used by phylogenetically distant luminous organisms [3]. At least 12 distinct types of luciferases, *sensu lato,* have been described [3].

When the photogenic reaction occurs, the generated light is defined by a color that can differ depending on luciferin, but also on the amino acid sequence and structure of the luciferase [4–7]. In some bioluminescent organisms, light emission involves Bioluminescence Resonance Energy Transfer (BRET), a nonradiative process where a bioluminescent donor (luciferase) transfers energy to a fluorescent acceptor (fluorescent protein) [4]. The fluorescent protein absorbs photons emitted by the bioluminescence reaction and becomes excited. It then re-emits light at longer wavelengths before returning to the ground state. For example, the Green Fluorescent Protein (GFP) emits green light after absorbing blue light primarily emitted by the luciferin-luciferase reaction [1,5].

Luminous anthozoans are distributed among four orders: Actiniaria (including one luminous families), Zoantharia (including two luminous families), Malacalcyonacea (including one luminous family), and Scleralcyonacea (**Table S1** ; [8]). Scleralcyonacea include 18 luminous families mainly spread into the Pennatuloidea superfamily (**Table S1** ; [8]). The bioluminescence system of *Renilla reniformis* (Pallas, 1766) (Pennatuloidea) has been extensively characterized. It involves the most widespread luciferin in marine environments, the coelenterazine (6-(4-hydroxyphenyl)-2-[(4-hydroxyphenyl)methyl]-8-(phenylmethyl)-7H-imidazo[1,2-a]pyrazin-3-one), and a coelenterazine-dependent luciferase (*R*Luc-type) [9–11,14,16]. This luciferase is homologous to bacterial haloalkane dehalogenases, assumed to have been horizontally transferred during evolution [3]. In addition, more recent descriptions of similar bioluminescent molecular components in different pennatulaceans are sparse [9–17] (**Table S1** ). Bessho-Uehara et al. (2020) demonstrated the widespread occurrence of coelenterazine-based *R*Luc-type luciferase across deep-sea pennatulacean species, such as *Distichoptilum gracile* (Verrill, 1882), *Umbellula* sp., *Pennatula* sp., and *Funiculina* sp. [13]. In addition, *R*Luc-type luciferases have also been identified in phylogenetically distant organisms such as echinoderms (*Amphiura filiformis* (Müller, 1776) [18]), and potentially in tunicates (*Pyrosoma atlanticum* (Péron, 1804) [19]), however, this has recently been called into question [20]. Luminescence-associated molecules, such as coelenterazine-binding proteins (CBPs) and GFPs have also been identified in sea pansy and other pennatulaceans [16,21–26]. Additionally, the calcium ion has been demonstrated to be indirectly involved in light production in *R. reniformis* and *Veretillum cynomorium* (Pallas, 1766) [27,28]. CBP relies on calcium ions as cofactors to release luciferin, enabling light production in *R. reniformis*[29].

Light production in sea pens occurs within specific endodermal cells (*i.e.,* photocytes), depending on the species, in the tissues of autozooid and siphonozooid polyps [30–32]. These photocytes often exhibit green autofluorescence [33,34]. To date, bioluminescent pennatulaceans display nervous catecholaminergic control of light emission, with adrenaline- and noradrenaline-triggered waves of light flashes [35,36]. Although no experimental behavioral evidence has been obtained, the ecological function of pennatulacean luminescence may serve as an aposematic signal, avoiding predation through misdirection or burglar alarm effects [2,37]. The current data and gaps in the anthozoan bioluminescence data are presented in **Table S1**.

Although the *Renilla* bioluminescence system has been extensively characterized, the molecular mechanisms and components involved in the bioluminescence of other sea pen species remain largely unexplored. This study aimed to explore the unique bioluminescent systems of three sea pen species, the pennatulid *Pennatula phosphorea* (Linnaeus, 1758), funiculinid *Funiculina quadrangularis* (Pallas, 1766), and anthoptilid *Anthoptilum murrayi* (Kölliker, 1880), focusing on their biochemical and molecular aspects. A multidisciplinary study using luminometric assays, transcriptome and phylogenetic analyses, 3D protein modeling, and immunodetection revealed the basis of the bioluminescent system of these soft coral sea pens. Cross-reaction luminometric results highlight the involvement of a *R*Luc-like luciferase in *P. phosphorea* and *F. quadrangularis*, both shallow water species, with coelenterazine as the substrate for all three bioluminescent systems. Besides, *P. phosphorea* and *A. murrayi de novo* transcriptome analyses corroborate (*i*) the homology of both retrieved luciferase sequences with the *Renilla* luciferase, (*ii*) the *in silico* presence of multiple CBPs-like and GFPs-like proteins homologous to the functionally characterized references proteins. The luciferase expression sites within the autozooids and siphonozooids of *P. phosphorea* and autozooids of *F. quadrangularis* are described. The presence of GFP was also strongly suggested in *P. phosphorea* through comparative microscopy. Finally, the pharmacological involvement of calcium in the light emission process of *P. phosphorea* and *F. quadrangularis* is confirmed.

## Materials and methods

### Specimen collection

In July 2022 and 2023, common sea pens (*Pennatula phosphorea*, n = 30), were collected from the Gullmarsfjord, Sweden, using a small dredge with a 1 m aperture at a depth of 35–40 m. Likewise, tall sea pens (*Funiculina quadrangularis*, n = 11), were sampled in July 2023 using the same method at a depth of 40 m. Animals were transported to the Kristineberg Marine Research Station (Fiskebäckskil, Sweden) and maintained under dark condition in a continuous flow of fresh deep-sea water pumped from the adjacent fjord.

Specimens of Murrays’ sea pens (*Anthoptilum murrayi*, n = 3) were collected during the DEEP-OCEAN expedition off the southeast coast of Brazil aboard the R/V Alpha Crucis. The collection was carried out using a demersal trawl net with 19 m lower rope, mesh sizes of 100 mm in the body and wings, and 25 mm in the codend, at an average depth of 1,500 m. Collection permits were issued by the *Instituto Chico Mendes de Conservação da Biodiversidade* (SISBIO permits #28054-4, 82624-1) and the *Secretaria da Comissão Interministerial para Recursos do Mar da Marinha do Brasil* (Portaria No. 223). After hoisting the net, all collected organisms were sorted, and *A. murrayi* specimens were promptly frozen in liquid nitrogen. These samples were stored at -80 °C at the Oceanographic Institute of the University of São Paulo (São Paulo, Brazil).

### Dissection & sample preparation

Specimens of *P. phosphorea* and *F. quadrangularis* were anesthetized by immersion in 3.5% MgCl_2_ solution in sea water for 30 minutes [36]. For each tested specimen of *P. phosphorea*, (*i*) the pinnules were dissected and weighed, and (*ii*) the rachis was divided into three equivalent portions and weighed. For specimens of *F. quadrangularis*, the rachis bearing the polyps was cut into 3 cm-long segments. Sea pen pinnules and rachises were either used directly for biochemical assays or rinsed for 3 h in fresh, running deep-sea water for pharmacological calcium assays. Other specimens were fixed in 4% paraformaldehyde (PFA) in phosphate buffer saline (PBS, 123 mM NaCl, 2.6 mM KCl, 12.6 mM Na_2_HPO_4_, 1.7 mM KH_2_PO_4_, pH 7.4) for subsequent immunodetection techniques.

Due to the challenges of collecting organisms from great depths (e.g., trawling damage), maintaining the survival of *A. murrayi* was not feasible. Consequently, all measurements were conducted on frozen samples.

### In vivo *and* in vitro *light emission spectrum of* Anthoptilum murrayi

The bioluminescence spectrum of *A. murrayi* was obtained using frozen coral pieces thawed and hydrated with deionized water. The spectra were recorded using a cooled CCD camera (LumiF SpectroCapture AB-1850). Measurements were conducted in triplicate, with an exposure time of 2 min at room temperature.

Similarly, the emission spectrum of the recombinant *A. murrayi* luciferase was recorded on the same equipment over a 60-second interval. Prior to this, the candidate *A. murrayi* luciferase coding sequence (obtained based on the method “*De novo transcriptome analyses for Pennatula phosphorea and Anthoptilum murrayi”* section) was optimized for *E. coli* expression and cloned into the pET-28a vector (Biomatik) between the BamHI and HindIII restriction sites. The N-terminal His-tagged luciferase was expressed in *E. coli* BL21 (DE3) cells, induced with 11mM IPTG at 15°C overnight. Cells were harvested and lysed in 50 mM sodium phosphate buffer (pH 7.4) containing lysozyme. The lysate was then purified using Ni-NTA affinity chromatography with an imidazole gradient (100–500 mM). Final luciferase preparations were pooled, dialyzed against 50 mM sodium phosphate buffer (pH 7.4), screened for light emission activity, and concentrated using a 3-kDa cutoff filter (Vivaspin 500, GE Healthcare). Unless otherwise stated, protein concentrations were measured using a Qubit 3.0 fluorometer (Invitrogen). The luminescent reaction was then conducted under conditions identical to the standard light emission assay. Specifically, 100 µL of solution containing *A. murrayi* luciferase was diluted with 3971µL of 501mM phosphate buffer (pH 7.4). The reaction was then initiated by adding 2 µL of native coelenterazine, bringing the total volume to 500 µL and yielding a final coelenterazine concentration of 6 µM.

### Luminescent system biochemistry

Eleven specimens of *P. phosphorea* were used for biochemical analysis. Tests were performed for the pinnules (n = 40), rachises (n = 10), and peduncles (n = 10). Pinnule locations along the axis were observed and classified as upper, middle, and lower, depending on the attachment position on the rachis, as in Duchatelet et al., 2023 [36]. Five specimens of *F. quadrangularis* were used immediately after dissection for biochemical assays. Measurements were performed independently of the polyp-bearing rachis location among the colonies (n = 10). Deep-frozen tissues from two *A. murrayi* specimens were used for this analysis. For the first two species, the central rigid calcified axis of the rachis was removed before starting experiments.

Light emission measurements were performed in a dark room using an FB12 tube luminometer (Tirtertek-Berthold, Pforzheim, Germany) calibrated with a standard 470 nm light source (Beta light, Saunders Technology, Hayes, UK). Light responses were recorded using FB12-Sirius PC Software (Tirtertek-Berthold). Light emission was characterized as follows: (*i*) maximum light intensity (Lmax), expressed in megaquanta per second (10^9^ q s^−1^), and (*ii*) total amount of light emitted (Ltot) over 3 min, expressed in megaquanta. All data were standardized per unit mass (g).

### *Luciferase and coelenterazine assays in* Pennatula phosphorea, Anthoptilum murrayi and Funiculina quadrangularis

For the luciferase assay, *P. phosphorea* pinnule and rachis and *F. quadrangularis* rachis were placed in 200 μl of Tris buffer (20 mM Tris, 0.5 mM NaCl; pH 7.4) and crushed with mortar and pestle until a homogenized extract was obtained; 20 and 40 μl of the extract was diluted in 180 and 160 μl Tris buffer, respectively. The diluted *P. phosphorea* and *F. quadrangularis* luciferase solutions were injected into two different tubes filled with 5 μl of a 1/200 stock solution of coelenterazine (Prolume Ltd., Pinetop, AZ, USA) in cold methanol (1OD at 430 nm) diluted in 195 μl of Tris buffer. Two measurements of Lmax were recorded and averaged to calculate the maximal light decay rate corresponding to luciferase activity expressed in 10^9^ q s^−1^ g^−1^ [1].

For coelenterazine detection, *P. phosphorea* pinnules, rachises, and *F. quadrangularis* rachises were placed in 200 μl of cold argon-saturated methanol and crushed using a mortar and pestle. Then, 5 μl of the methanolic extract was injected into a tube filled with 195 μl of Tris buffer and placed in a luminometer. Afterwards, 200 μl of *Renilla* luciferase solution constituted 4 μl of *Renilla* luciferase (Prolume Ltd., working dilution of 0.2 g l^−1^ in a Tris-HCl buffer 10 mM, NaCl 0.5 M, BSA 1%; pH 7.4) and 196 μl of Tris buffer was injected into the luminometer tube. The Ltot was recorded and used to calculate the amount of coelenterazine contained in a gram of pinnule (ng g^−1^), assuming that 1 ng of pure coelenterazine coupled with *Renilla* luciferase emits 2.52 × 10^11^ photons [1].

*A. murrayi* tissues were processed using a Potter-Elvehjem homogenizer in 2 mL of Tris-HCl buffer (50 mM, pH 8.0). Following homogenization, the mixture was centrifuged at 15,000 × g for 10 min at 4°C and the pellet was discarded. The supernatant was used for the light emission assays. For each assay, the mixture comprised 100 µL of supernatant, 397 µL of Tris-HCl buffer (50 mM, pH 7.4), and 3 µL of coelenterazine (Prolume Ltd., Pinetop, AZ, USA), to achieve a final volume of 500 µL with a final concentration of 6 µM.

Control measurements were performed using coelenterazine in the presence of the buffer, and the emitted light remained around 1,000 RLU. This value was subtracted from the light emitted by the enzymatic assays for standardization purposes. All luminescence assays were conducted in triplicate, and the final result represents the average of the triplicates.

### *Long-term light monitoring of coelenterazine production in* Pennatula phosphorea

The *P. phosphorea* specimens (n = 12) were maintained in tanks filled with circulating artificial seawater (ASW; 400 mM NaCl, 9.6 mM KCl, 52.3 mM MgCl_2_, 9.9 mM CaCl_2_, 27.7 mM Na_2_SO_4_, 20 mM Tris; pH 8.2) at temperatures following natural temperature variations encountered in the native fjord (https://www.weather.mi.gu.se/kristineberg/en/data.shtml) with a 12-12 hours photoperiod. Sea pens were fed weekly with REEF LIFE Plancto (Aqua Medic, Germany), a food without any coelenterazine trace. After six and 12 months, potassium chloride (KCl) depolarization, coelenterazine content, and luciferase activity assays were performed on eight and four specimens, respectively, with two replicates per specimen. Luciferase and coelenterazine assays were performed as previously described. Total depolarization through KCl application allows for rapid estimation of the luminous ability of specimens. For the KCl experiments, luminescence induction was performed on the pinnule and rachis portions placed in a tube luminometer filled with 500 µL of ASW. Then, the light emission was triggered with the addition of 500 µL of a KCl solution (400 mM KCl, 52.3 mM MgCl_2_, 9.9 mM CaCl_2_, 27.7 mM Na_2_SO_4_, 20 mM Tris; pH 8.2), and Ltot was recorded over 3 minutes. The recorded luciferase activity, luciferin content, and KCl response were compared with wild-caught measurements.

### De novo *transcriptome analyses for* Pennatula phosphorea *and* Anthoptilum murrayi

The pinnule, rachis, and peduncle tissues of a single *P. phosphorea* specimen were dissected and directly immersed in a permeabilizing RNAlater-Ice (Life Technologies) solution overnight at -20°C, following the manufacturer’s protocol. Subsequently, the samples were stored at -80°C and processed for RNA extraction. Total RNA was extracted using the TRIzol reagent. The quality of the RNA extracts (RIN value, fragment length distribution, and 28S/18S ratio) and their concentrations were assessed using an Agilent 2100 bioanalyzer. The BGI company (Beijing Genomics Institute, Hong Kong) performed cDNA library preparation and sequencing using a procedure similar to that previously described [38–40]. High-throughput sequencing was conducted using the BGISEQ-500 platform to generate 100 bp paired-end reads. To exclude low-quality sequences, the raw reads were filtered by removing (*i*) reads with more than 20% of the qualities of the base lower than 10, (*ii*) reads only containing the adaptor sequence, and (*iii*) reads containing more than 5% of unknown nucleotide “N.” Quality control of the reads was performed using FastQC software [41]. For *P. phosphorea,* a reference *de novo* transcriptome assembly was then created from the remaining clean reads obtained from the pinnule, rachis, and peduncle tissues using Trinity software [42] (Trinity-v2.5.1; min_contig_length 150, CPU 8, min_kmer_cov 3, min_glue 3, SS_lib_type RF, bfly_opts’-V 5, edge-thr=0.1, stderr’). TGICL software was then used to reduce transcriptome redundancy by assembling the contigs into a single set of longer, non-redundant, and more complete consensus unigenes [43] (Tgicl-v2.5.1; -l 40 -c 10 -v 25 -O ’-repeat_stringency 0.95 -minmatch 35 -minscore 35’). Unigenes, defined as non-redundant assembled sequences obtained from assembly and/or clustering [44], can form clusters in which the similarity among overlapping sequences is greater than 70% or singletons that are unique unigenes. For all transcriptomes, unigene expression was evaluated using the “Fragments per kilobase of the transcript, per million fragments sequenced” (FPKM) method [38–40,44]. To obtain annotation for transcriptomes, unigenes were aligned to NCBI Nucleotide (NT), NCBI protein (NR), EuKariotic Orthologous groups (KOG), Kyoto Encyclopedia of Genes and Genomes (KEGG), and UniProtKB/Swiss-Prot databases using Blastn, Blastx [45], and Diamond [46]. Blast2GO [47] with NR annotation results was used to obtain Gene Ontology annotations according to molecular function, biological process, and cellular component ontologies. InterPro [48] was also used to annotate unigenes based on functional analyses of protein sequences, clustering them into families and predicting the presence of domains or essential amino acid residues. The candidate coding area among the unigenes was assessed using Transdecoder (https://transdecoder.github.io).

A similar procedure was used for the specimen of *A. murrayi*. Whole sea pen tissues (pinnule and peduncle) were dissected, frozen in liquid nitrogen, and stored at -80°C. The total RNA from the sea pen was extracted using the RNeasy Plant Mini Kit (QIAGEN), following the manufacturer’s instructions. RNA samples were treated with DNase I (Invitrogen) to remove potential genomic DNA contamination. Subsequently, to remove impurities and residual DNase I reaction remnants, they underwent a column cleanup process, following the instructions of the QIAGEN Kit. The concentrations of RNA samples were estimated by fluorescence using a Qubit fluorometer (Thermo Fisher Scientific). The integrity of the RNA samples was confirmed by agarose gel electrophoresis (1%) stained with SYBR Safe (Invitrogen), and the RNA was dried at 30°C for 1 h in speed vac mode V-AQ.

The quality of RNA extracts (RIN value, fragment length distribution, and 28S/18S ratio) and their concentration were assessed by RNA concentrations were evaluated using an Agilent 2100 bioanalyzer. After QC, mRNA was enriched using oligo(dT) beads. First, mRNA was randomly fragmented by adding a fragmentation buffer. Then, the cDNA was synthesized using an mRNA template and random hexamer primer, after which a custom second-strand synthesis buffer (Illumina), dNTPs, RNase H, and DNA polymerase I were added to initiate second-strand synthesis. After a series of terminal repairs, A ligation, and sequencing adaptor ligation, the double-stranded cDNA library was completed through size selection and PCR enrichment. The GenOne Biotechnologies company (Brazil) performed cDNA library preparation and sequencing (150 bp paired-end reads). To exclude low-quality sequences, the raw reads were filtered by removing (*i*) reads with more than 50% of the qualities of the base lower than 5, (*ii*) reads only containing the adaptor sequence, and (*iii*) reads containing more than 10% of unknown nucleotide “N.”

RNA sequencing was performed with v4 21×11001bp reads on the HiSeq2500 platform by NGS Soluções Genômicas in Piracicaba, São Paulo. The reference *de novo* transcriptome assembly was generated using Trinity v2.13.2. To differentiate among identified isoforms, transcript abundance was analyzed using the *align_and_estimate_abundance.pl* script from the Trinity software package [49].

Transcriptome completeness was evaluated using BUSCO v4 (Benchmarking Universal Single-Copy Orthologs), a bioinformatics tool that determines the proportion of single-copy orthologs present in the dataset Metazoa_odb9. The BUSCO analysis and visualization of the results were conducted on the Galaxy platform (https://usegalaxy.org).

### Light emission process-related protein sequence analyses

Potential transcripts of interest were chosen using NCBI online tools according to potential phylogenetic homologies to identify genes involved in the light production process, such as luciferases, green fluorescent proteins, and coelenterazine-binding proteins. These “light emission process-related genes” were searched within the newly generated *P. phosphorea* transcriptome using tBLASTn analysis (1 hit, E-value < 1e^-20^). All retrieved unigenes were individually reciprocally searched in the NCBI NR database (Reciprocal BLASTx; 1hit, E-value < 1e^-20^). BLAST hits with significant E-values strongly support homologous proteins. *In silico* translation (ExPASy translate tool, http://expasy.org/tools/dna.html) was performed on the sequences retrieved from the *P. phosphorea* and *A. murrayi* transcriptomes for all putative candidates. Multiple alignments were performed for each predicted protein with their respective homologous “light emission process-related protein” retrieved from the NCBI online tool in other species using Geneious software [50]. Sequence alignments have enabled the identification of luciferase characteristic features, such as catalytic triads [51].

To validate the *P. phosphorea* luciferase retrieved sequence, primers were designed based on *Renilla muelleri* luciferase mRNA (AY015988.1) to amplify the hypothetical luciferase sequence using Primer Blast software (**Table S2** ). *P. phosphorea* pinnules were collected and lysed in 300 µL of 50 mM NaOH for 30 min at 95°C with agitation at 800 rpm. The pH was adjusted by adding 60 µL 500 mM Tris-HCl at pH 8. For PCR amplification, the reaction mix contained 1U of Expand Long Template (Roche) with the provided buffer, 400 µM dNTP (#R0191, Life Technologies), and 250 nM of each primer. Amplification was performed as follows: 95°C for 5 min; 35 cycles of denaturation at 95°C for 30 s, hybridization from 52 °C to 55°C for 15 s, and elongation at 68°C for 45–80 s. The final cycle was completed with a final elongation step at 68°C for 7 min. Primer pairs F1-R1, F2-R2, and F8-R2 resulted in amplifications visualized through electrophoresis and ethidium bromide incorporation. Genomic DNA was purified using the Qiagen PCR Purification Kit according to the manufacturer’s instructions and sequenced by Microsynth.

### Phylogenetic analyses

The predicted sequences of the luciferases (LUC), GFP, and CBPs of *P. phosphorea* and *A. murrayi* were placed into an anthozoan-focused phylogenetic context using maximum likelihood phylogenetic reconstruction [52]. *Renilla*-type luciferase sequences from selected metazoans were collected from public databases based on literature [13,17,18]. GFP sequences from cnidarians were collected from public databases using reference literature [53–56]. CBP sequences from anthozoans were collected from public databases based on literature [57]. Multiple alignments of all sequences were performed using the MAFFT algorithm implemented in the Geneious software and trimmed using TrimAL software [58]. Maximum likelihood phylogenetic analyses were performed using IQ-tree software [59]. Before the analyses, ModelTest [60] was used to select the best-fit evolution model. Trees were edited using the iTOL web tool.

### Structural modeling

AlphaFold models of *P. phosphorea* and *A*. *murrayi* LUC, GFP, and CBP proteins were obtained from the AlphaFold web server (https://alphafoldserver.com/). The structural models were visualized using PyMOL 2.6 (https://pymol.org/). Structural pairwise alignments and calculations were performed using the Dali server (http://ekhidna2.biocenter.helsinki.fi/dali/).

### Luciferase and green fluorescent protein immunodetection

Commercial antibodies against *Renilla* luciferase (GTX125851, Genetex) were used to confirm the presence of a *Renilla*-like luciferase. Proteins were extracted from frozen pinnules, rachis, and peduncle samples. Each sample was homogenized on ice in 1000 μl of 2% Triton X-100 in phosphate buffer saline (PBS: 10 mM Tris, pH 7,5; 1 mM EDTA, pH 8,0; 100 mM NaCl) supplemented with protease inhibitors (complete–Mini tablets, Roche). The extract was sonicated and centrifuged at 800 g for 15 min. The supernatant was then collected. The protein concentration in each extract was measured using a PierceTM BCA Protein Assay Kit (Thermo Scientific). Laemmli buffer (Bio-Rad) and β-mercaptoethanol (βMSH, Bio-Rad) were added to each protein extract and the proteins were electrophoretically separated at 200 V for 35 min on 12% SDS-PAGE gels. The separated proteins were electroblotted onto nitrocellulose membranes. The membrane was incubated overnight with the primary anti-*Renilla* luciferase antibody and the secondary antibody (ECL HRP-conjugated anti-rabbit antibody, Life Sciences, NA934VS, lot number 4837492) for 1 h. Antibody detection was performed using the reagents of the detection kit (HRP Perkin-Elmer, NEL 104) following the manufacturer’s instructions. The dilution for the primary antibody was 1:2000.

The same primary antibody was used to immunolocalize luciferases within the *P. phosphorea* pinnules, rachis, peduncle tissues, and *F. quadrangularis* polyp tissue. For *whole-mount* immunofluorescence, the dissected samples were blocked with PBS containing 2% Triton X-100 and 6% BSA (Amresco). Samples were then incubated for 48 h with either the anti-*Renilla* luciferase antibody diluted 1:200 in PBS containing 1% Triton X-100, 0.01% NaN_3_, and 6% BSA. Visualization of the luciferase signal was performed after 24 h of incubation of the samples at RT in the dark with a fluorescent dye-labeled secondary antibody (Goat Anti-Rabbit, Alexa Fluor 594, Life Technologies Limited) diluted 1:500 in PBS containing 1% Triton X-100, 0.01% NaN_3_, and 6% BSA. Samples were mounted (Mowiol 4–88, Sigma) and examined using an epifluorescence microscope (Axio Observer, Zeiss, Oberkochen, Germany) equipped with Zen microscopy software (Zeiss, Oberkochen, Germany). Control sections were incubated in PBS containing 1% Triton X-100, 0.01% NaN_3_, and 6% bovine serum albumin (BSA) with no primary antibodies.

### Calcium assays

Following the identification of the key proteins involved in the luminescence system, we assessed the role of calcium, another crucial element in the bioluminescence mechanism. Different pharmacological tests were used to investigate the role of calcium in the light-emission process of *P. phosphorea* and *F. quadrangularis* (n = 6 for each species). First, the calcium concentration was tested using three artificial seawater (ASW) solutions with different calcium concentrations (0, 10, and 20 mM CaCl_2_). To remove any traces of Ca^2+^ ions in the 0 mM CaCl_2_ -ASW solution, calcium chelators EGTA (4100, Merck) and BAPTA (14513, Merck) were added to the solution at 10^-5^ M final concentrations. Secondly, the effect of a calcium ionophore (A23187; C7522, Merck) was tested to highlight the potential involvement of calcium storage in the light emission process. Third, the involvement of calcium was tested in the presence of a previously determined triggering agent of light emission in sea pens, adrenaline at 10^-5^ M. Finally, calcium involvement was tested on the effect of the potassium chloride depolarization solution usually employed to trigger the maximum light production in a bioluminescent species.

After rinsing for 3 h, the pinnules of *P. phosphorea* and polyp-bearing rachises of *F. quadrangularis* were placed in luminometer tubes and subjected to various treatments (**Table S3** ). Before the experiments, each sample was pre-incubated for 15 min in ASW devoid of calcium (0 mM CaCl_2_). Data were recorded for 15 minutes using an FB12 tube luminometer (Tirtertek-Berthold, Pforzheim, Germany). Light emission was defined as the total amount of light emitted (L_tot_) over 15 min, and was expressed in megaquanta. All data were standardized per unit mass (g).

### Statistical analysis

All statistical analyses were performed with R Studio (version 2023.03.1 + 446, 2022, Posit Software, USA). Variance normality and equality were tested using the Shapiro–Wilk test and Levene’s test, respectively. When these parametric assumptions were met, Student’s t-test and ANOVA coupled with Tukey’s test were used to perform single or multiple comparisons between groups. When log transformation did not provide normality and homoscedasticity, the nonparametric Wilcoxon test and Kruskal–Wallis test coupled with the Wilcoxon rank-sum test were used to assess whether significant differences were present between the two groups or multiple groups. Differences were considered significant at a minimum *p-value* of < 0.05. Values are graphically illustrated as the mean and standard error of the mean (s.e.m).

## Results

### Pennatula phosphorea, Funiculina quadrangularis*, and* Anthoptilum murrayi *emit light using a coelenterazine-dependent luciferase*

Biochemical assays performed on *P. phosphorea, F. quadrangularis*, and *A. murrayi* (**Figures 1A-F**) demonstrated a similar response pattern. The cross-reactivity of the extracts with the commercial *Renilla* luciferase highlights the involvement of coelenterazine in the bioluminescent systems of *P. phosphorea, F. quadrangularis*, and *A. murrayi* (**Figure 1**). For *P. phosphorea*, the measured amounts of coelenterazine present no statistical differences between the pinnule areas (ANOVA, *p-value* lower-middle = 0.8122; *p-value* lower-upper = 0.9054; *p-value* upper-middle = 0.7014) (**Figure S1**). Similarly, no significant differences were observed between the rachis portions (ANOVA, *p-value* lower-middle = 0.5905; *p-value* lower-upper = 0.8274; *p-value* upper-middle = 0.4634) (**Figure S1**). The mean coelenterazine content per pinnule and rachis were 131.15 ± 29.06 and 70.00 ± 16.79 ng g^-1^, respectively. Peduncle presents an almost negligible mean value of 0.21 ± 0.01 ng g^-1^. Comparatively, the mean coelenterazine content of *F. quadrangularis* polyp-bearing rachis was 3.79 ± 1.20 ng g^-1^.

**Figure 1.**
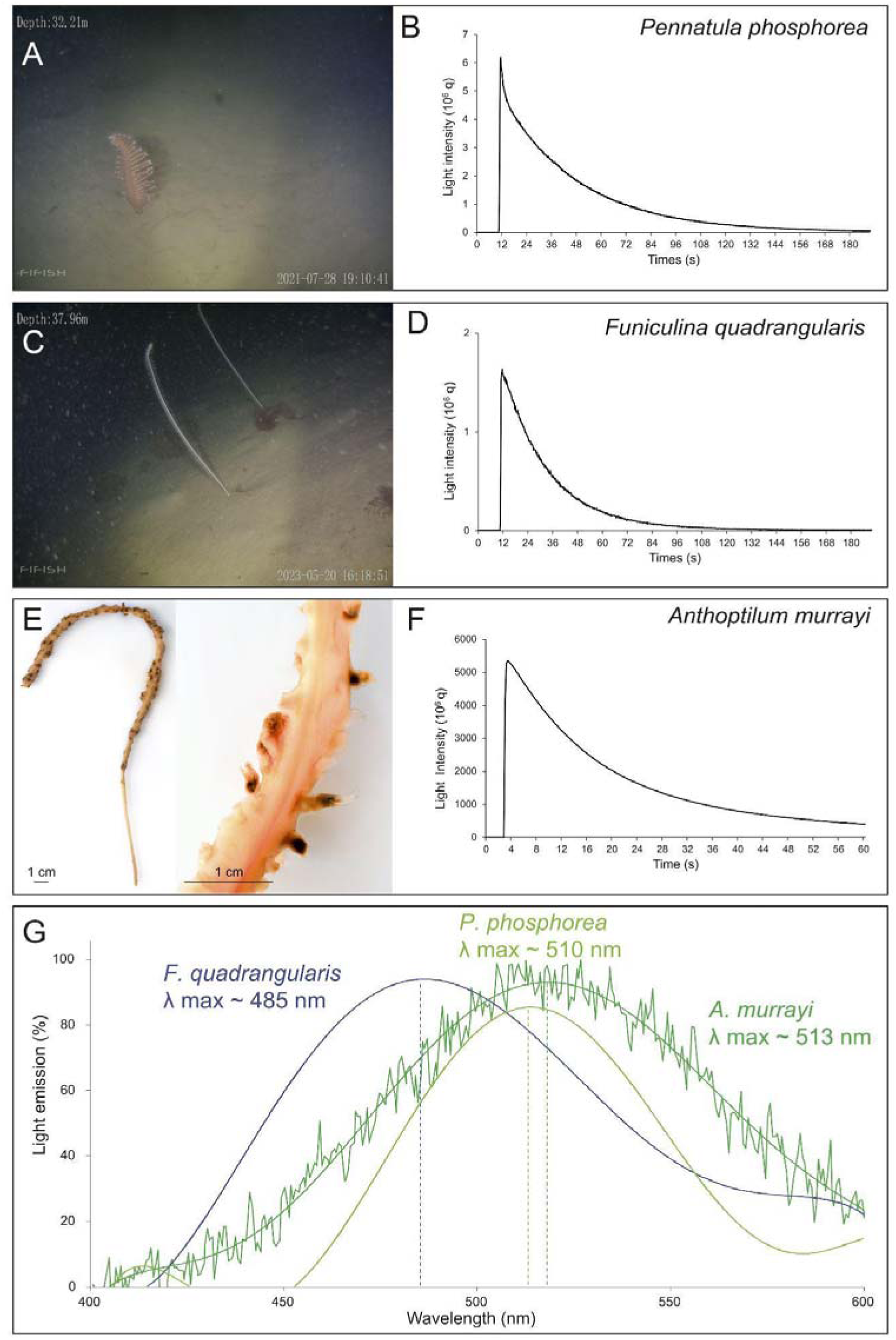
Biochemistry assays. (A) *In situ* images of *Pennatula phosphorea* and (B) typical coelenterazine assay curve for the pinnules. (C) *In situ* images of *Funiculina quadrangularis* and (D) typical coelenterazine assay curve for the 3 cm-long polyp-bearing rachis. (E) Images of *Anthoptilum murrayi* with a zoom on the polyp on the rachis and (F) typical coelenterazine assay curve for the whole specimen. (G) *Anthoptilum murrayi in vivo* luminescence spectrum compared with the retrieved spectrum of two other species (*P. phosphorea* [61] and *F. quadrangularis* [13]). Pictures (A and C) were provided by Fredrik Gröndahl.

In parallel, a typical coelenterazine assay curve was observed for all three species (**Figures 1B, D, and F)**. Cross-reaction assays using synthetic coelenterazine to measure potential luciferase activity confirmed the involvement of a coelenterazine-based luciferase system in *P. phosphorea* and *F. quadrangularis*. For *P. phosphorea*, luciferase activities in the pinnules show no statistically significant differences across the pinnule areas (ANOVA, *p-value* lower-middle = 0.4963; *p-value* lower-upper = 0.2742; *p-value* upper-middle = 0.6251), with a mean Lmax value of 129.5 ± 22.1 10^9^ q g^-1^ s^-1^ (**Figure S1** ). Similarly, no statistically significant differences were observed in the Lmax values recorded between rachis portions (ANOVA, *p-value* lower-middle = 0.6360; *p-value* lower-upper = 0.4438; *p-value* upper-middle = 0.2548), with a mean Lmax value of 87.9 ± 16.9 10^9^ q g^-1^ s^-1^ (**Figure S1**). Finally, the peduncles tested exhibited a significantly lower mean Lmax value (0.6 ± 0.4 10^9^ q g^-1^ s^-1^) compared to the rachis (ANOVA, *p-value* = 0.0298). Comparatively, *F. quadrangularis* luciferase activity presents a mean Lmax value of 313.8 ± 99.2 10^9^ q g^-1^ s^-1^. To complete the data on the sea-pen light emission spectrum, the *in vivo* light emission spectrum of *A. murrayi* was measured and revealed a peak wavelength at 513 nm (**Figure 1G**).

### Pennatula phosphorea *maintains its bioluminescence ability after one year in captivity without an exogenous supply of coelenterazine*

Before the experiment, visual assessments of *P. phosphorea* luminescence were performed in the dark. These observations confirmed that the species maintained the ability to produce visible light even after six–12 months of captivity without any external sources of coelenterazine-containing food.

Measurements of luminescence parameters showed a general decrease in all three parameters for both the pinnules and the rachis (**Figure S2**). For the pinnules, the maximum light emission shows statistically significant differences upon KCl application (Kruskal-Wallis test, *p-value* = 1.8 10^-5^), with a mean Ltot value of 361.7 ± 34.6 10^9^ q g^-1^ s^-1^ after field collection. This value is statistically different from the mean value observed after six months of captivity (Wilcoxon sum-rank test, *p-value* = 1.5 10^-6^) but not from the value after 12 months (Wilcoxon sum-rank test, *p-value* = 0.16). The mean Ltot values after 6 and 12 months are 80.5 ± 21.8 and 223.8 ± 84.4 10^9^ q g^-1^ s^-1^, respectively. Coelenterazine content also presented statistically significant differences (Kruskal-Wallis test, *p-value* = 1.3 10^-5^). The mean coelenterazine content is already drastically reduced after 6 months (30.7 ± 4.5 ng g^-1^) but remains stable for one year (28.1 ± 4.4 ng g^-1^) (Wilcoxon sum-rank test, T0 *VS* T6: *p-value* = 0.0001; T0 *VS* T12: *p-value* = 0.0004), without statistical differences between the former two (Wilcoxon sum-rank test, *p-value* = 0.91) (**Figure S2**). Similarly, the luciferase activities decreased over the year and presented statistically significant differences (Kruskal-Wallis test, *p-value* = 0.01). Differences occur between the initial Lmax value (113.2 ± 12.7 10^9^ q g^-1^ s^-1^) and the mean Lmax values of 77.0 ± 16.2 and 43.0 ± 3.7 10^9^ q g^-1^ s^-1^ after 6 and 12 months, respectively (Wilcoxon sum-rank test, T0 *VS* T6: *p-value* = 0.383; T0 *VS* T12: *p-value* = 0.013; T6 *VS* T12: *p-value* = 0.383) (**Figure S2**). Similar observations occurred for the three parameters recorded on the rachis portions of *P. phosphorea* (**Figure S2**).

### *De novo transcriptomes of* Pennatula phosphorea *and* Anthoptilum murrayi

For *P. phosphorea*, a total of 47.27 million raw reads of 200 bp length were generated from the pinnule library, 47.27 million from the rachis library, and 41.96 million from the peduncle library. Data quality was assessed using FastQC software. Raw reads are available on the NCBI SRA database: *A. murrayi* (PRJNA1144931), *P. phosphorea* (PRJNA1152785). After low-quality reads filtering, the remaining high-quality reads (*i.e.,* 45.29 for the pinnule transcriptome, 45.53 for the rachis transcriptome, and 40.29 for the peduncle transcriptome) were used to assemble a reference transcriptome using the Trinity software. The obtained Trinity-predicted transcripts were clustered using TGICL to obtain the final unigenes.

In total, 49,510 unigenes (i.e., non-redundant unique sequences) were obtained with a total length of 72,928,901 bp. The average length was 1,473 bp and the N50 was 2,350 bp. Among the transcriptome data, 35,439 unigenes for the pinnule dataset, 33,778 unigenes for the rachis dataset, and 31,001 unigenes for the peduncle dataset were obtained, with a total of 49,510 different unigenes. The length distributions of the unigenes are shown in **Figure 2A** and the numerical data are summarized in **Tables S4** and **S5**.

**Figure 2.**
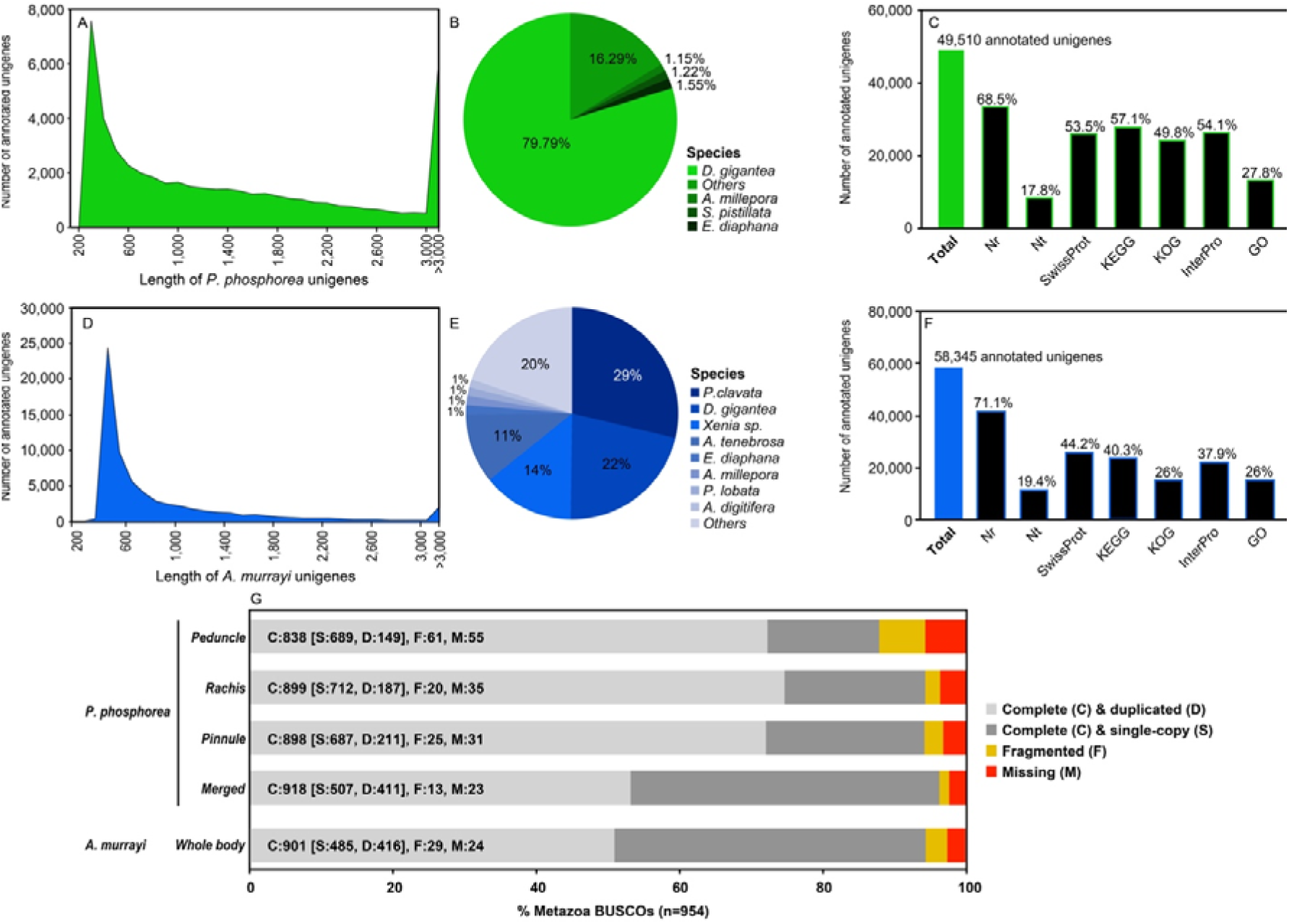
General description of the transcriptomic data. (A) The length distribution of *P. phosphorea* unigenes. (B) Taxonomic annotation of *the P. phosphorea* transcriptome. (C) Global annotation of *P. phosphorea* transcriptome. (D) Length distribution of *the A. murrayi* unigenes. (E) Taxonomic annotation of *A. murrayi* transcriptome. (F) Global annotation of *the A. murrayi* transcriptome. (G) Busco analyses.

Of the 46,258 predicted unigenes, 1,690 were found only in the pinnule transcriptome, 1,114 only in the rachis transcriptome, 1,035 in the peduncle transcriptome, and 36,934 in the peduncle transcriptome. For descriptive purposes, comparative gene expression analysis was performed by mapping FPKM values (*e.g.,* log_10_(FPKM value pinnule transcriptome) against log_10_ (FPKM value rachis transcriptome)) calculated for all predicted unigenes (**Table S6** ). However, it has to be clarified that transcriptome data have been generated for new gene discovery, not differential expression analyses, as no biological or technical replication was performed as a part of the study. The main species represented within the unigene annotation of the reference transcriptome was the anthozoan *Dendronephthya gigantea* (Verrill, 1864) (79%) (**Figure 2B** ). Of the 49,510 *P. phosphorea* unigenes present in the filtered reference transcriptome, 34,984 showed significant matches with the molecular databases: 33,896 to NR (68.5%, E-value < 1e^-5^), 8,818 to NT (17.8%), 26,473 to SwissProt (53.5%), 28,290 to KEGG (57.1%), 24,642 to KOG (49.8%), 26,790 to InterPro (54.1%), and 13,742 to GO (27.8%) (**Figure 2C**).

For *A. murrayi*, 66,425 unigenes were obtained, with a total length of 54,366,500 bp. The average mean length was 818 bp and the N50 was 1140 bp. The length distributions of the unigenes are shown in **Figure 2D** and the numerical data are summarized in **Tables S4** and **S5**. For descriptive purposes, gene expression analysis was performed by mapping the FPKM values (e.g., log_10_(FPKM value)) calculated for all predicted unigenes (**Table S6** ). The main represented species within the unigene annotation of the reference transcriptome was the anthozoan *Paramuricea clavata* (29%) (**Figure 2E** ). Among the 66,425 *A. murrayi* unigenes present in the filtered reference transcriptome, 58,345 were significantly matched to the molecular databases: 41,512 to NR (71.15%, E-value < 1e-10), 11,345 to NT (19.44%), 25,769 to SwissProt (44.16%), 23,549 to KEGG (40.36%), 15,178 to KOG (26.01%), 22,134 to InterPro (37.93%), and 15,178 to GO (26.01%) (**Figure 2F**).

Because of the lack of a reference genome in *P. phosphorea* and *A. murrayi*, and possibly the relatively short length of some unigene sequences, 29.4% and 71.6%, respectively, of the assembled sequences could not be matched to any known genes.

Based on Benchmarking Universal Single-Copy Orthologs (BUSCO) analysis, 96.23% of metazoa BUSCO genes were predicted to be complete in the merged *P. phosphorea* transcriptome. In parallel, 1.36% of BUSCO genes were fragmented and 2.41% were missing (**Figure 2G** ). Similar results were obtained for *A. murrayi*, with 94.44% complete BUSCO genes found, while 3.04% of BUSCO genes were fragmented, and 2.52% were missing (**Figure 2G**).

Analyses of unigene expression revealed a similar proportion of FPKM values in the three tissues (**Figure 3A, B** ). Comparatively, a large number of unigenes appeared to be more highly expressed in the pinnule and rachis than in the peduncle tissue (**Figure 3C**). The pinnule and rachis tissues appeared to be more similar in terms of the unigene expression profile. The “Molecular function” GO functional annotations for the 40 most expressed unigenes of each sample of *P. phosphorea* (**Figure 3D**) and the whole animal sample for *A. murrayi* (**Figure 3F** ) show a classical high expression of molecular actors involved in the cellular machinery and gene regulation. The GO functional annotations for the 40 unigenes with the most pronounced expression differences between tissues, including those highly expressed in one tissue relative to the other and those with lower expression levels, are shown in **Figure 3F**. We did not perform formal differential gene expression analysis, as no replication was performed. Calcium ion binding function (GO: 0005509) appeared to be predominantly expressed in pinnule and rachis tissues, compared to the peduncle (**Figure 3F**). Similarly, the pinnule tissue (containing the feeding polyp) presented a higher proportion of genes annotated as digestive enzymes (**Figure 3F**).

**Figure 3.**
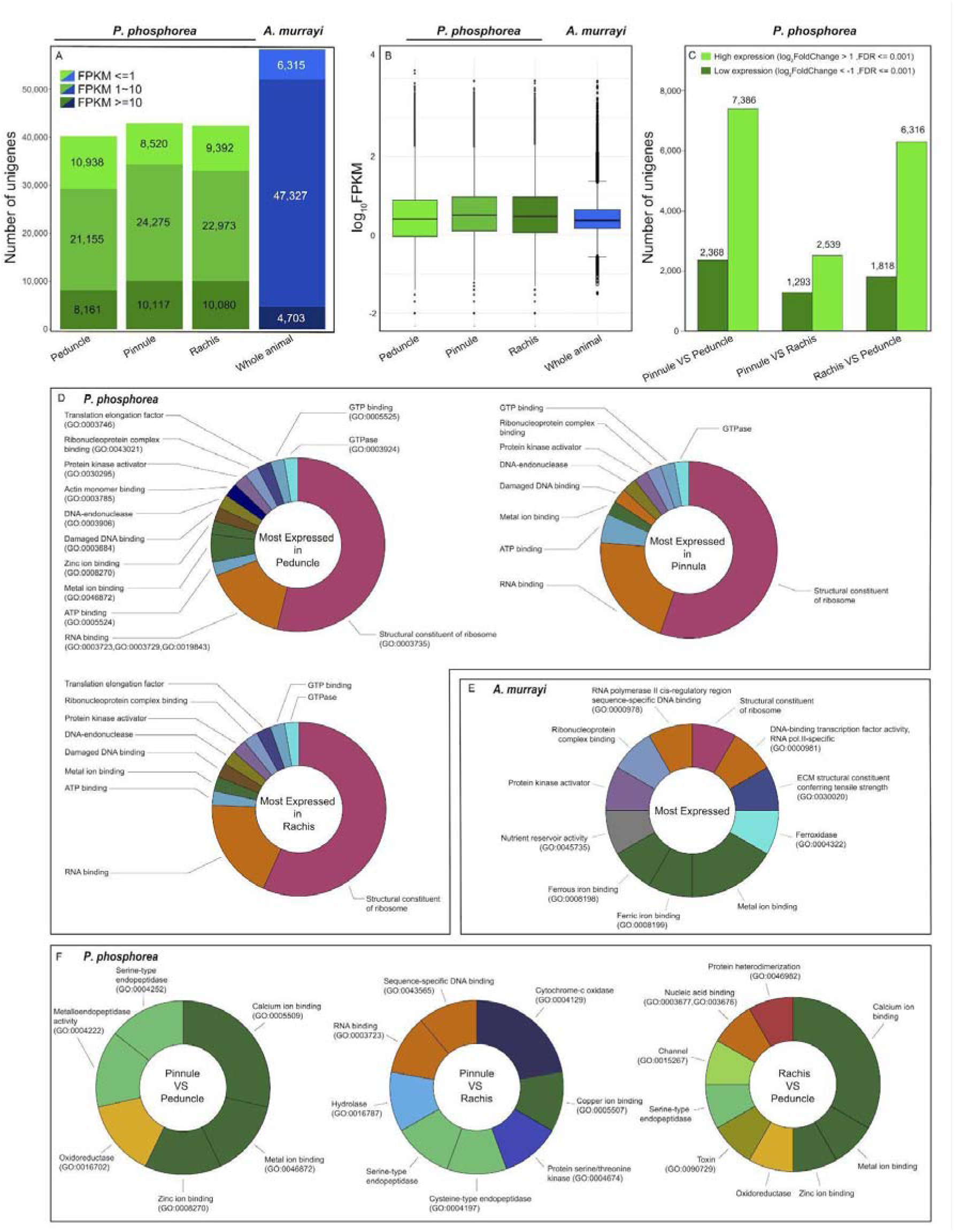
Global gene expression and gene ontology of *Pennatula phosphorea* and *Anthoptilum murrayi* transcriptomic data. (A) Distribution of FPKM expression values across *the P. phosphorea* and *A. murrayi* transcriptomes. (B) Gene expression distribution for each sample. (C) Proportions of unigenes with relatively high or low expression levels in *P. phosphorea* samples. Gene ontology distribution of the 40 most highly expressed unigenes for each sample of *P. phosphorea* (D) and *A. murrayi* (E). (F) Comparison of gene ontology repartition of the 40 most highly expressed unigenes in *P. phosphorea* samples.

### Expression of bioluminescence-related genes in Pennatula phosphorea and Anthoptilum murrayi

The *P. phosphorea* and *A. murrayi* transcriptomes contained sequences of several predicted luciferases, GFPs, and luciferin-binding proteins. The FPKM values retrieved from each transcriptome are shown in **Table S6**. Reciprocal BLAST analyses revealed that the sequences matched the luciferases, CBPs, and GFPs of Anthozoans.

Several *R*Luc-like enzymes were recovered from both investigated species (*A. murrayi* and *P. phosphorea*). In addition, sequence mining allowed us to recover additional *R*Luc-like sequences from the genomes of *Renilla muelleri* and *R. reniformis*. Phylogenetic analyses revealed three clades of *R*Luc-like enzymes in Pennatulacea (**Figure 4A** ). Clade A contains well-known luciferases from *R. muelleri* and *R. reniformis* and probable luciferase sequences from *A. murrayi, P. phosphorea,* and *Cavernularia obesa* (Valenciennes, 1850). Interestingly, clade A did not contain any sequences from the non-luminous species. Clade B, in comparison, includes several sequences from non-luminous species (*e.g., Pinnigorgia flava* (Nutting, 1910) and *Sinularia cruciata* (Tixier-Durivault, 1970)), in addition to several sequences from both our model species. Clade C also contained *the R* Luc-like enzymes retrieved from each transcriptome.

**Figure 4.**
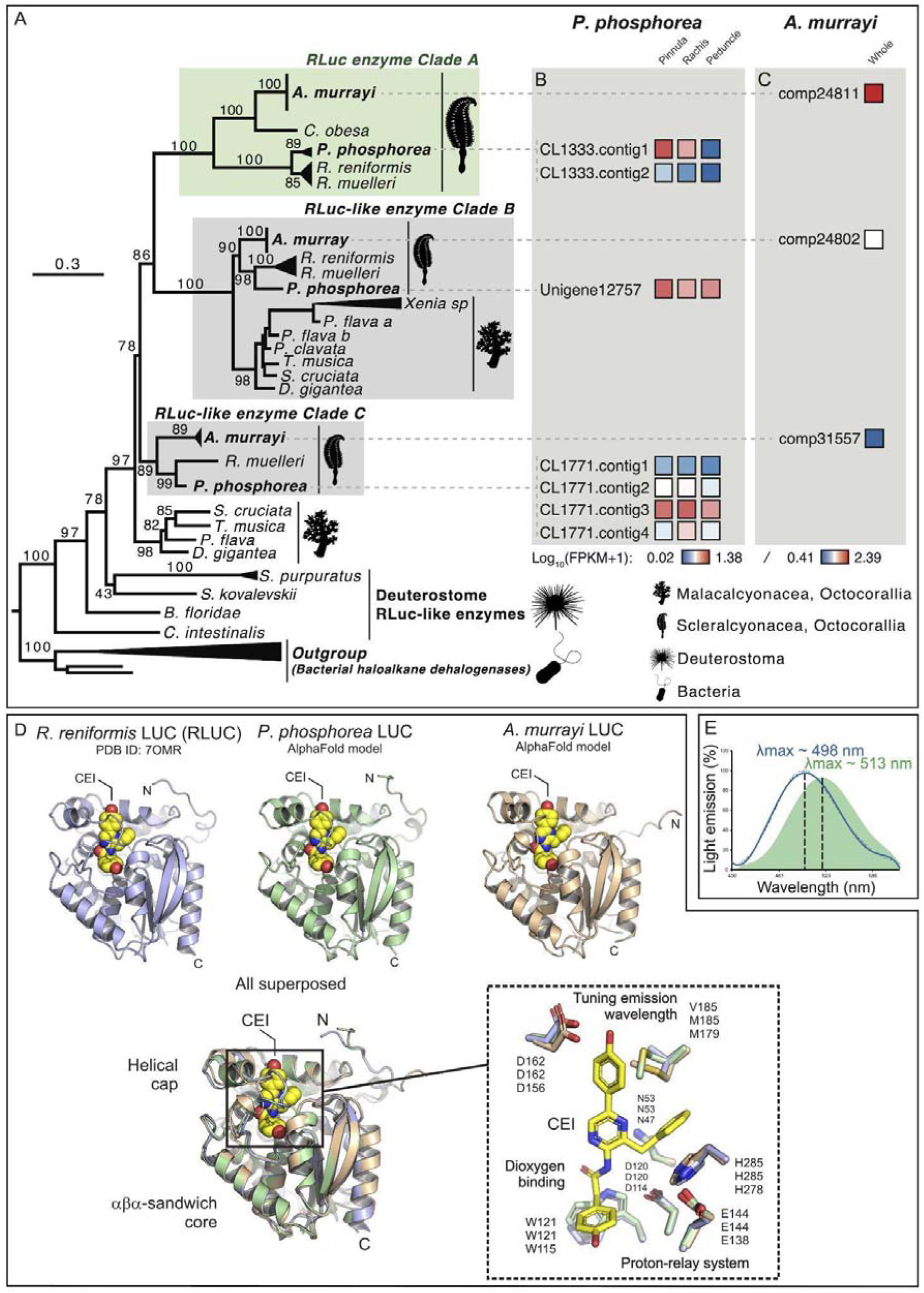
Phylogenetic tree of *R*Luc-like enzymes, including *P. phosphorea* and *A. murrayi* amino acid sequences. (A) Maximum likelihood tree based on the amino acid sequence alignment of *R*Luc-like enzymes. The tree was calculated by the IQ-tree software using the LG+I+G4 model of evolution. Numbers at the nodes indicate ultrafast bootstrap values based on 1000 replicates. The scale bar represents the percentage of amino acid substitutions per site. Bacterial haloalkane dehalogenase sequences were used to root the tree. (B) The expression level of each retrieved *R*Luc-like sequence in a distinct portion of *P. phosphorea*. (C) Expression level of each retrieved *R*Luc-like sequence in *the whole A. murrayi* specimen. (D) Structural comparison of AlphaFold models of *P. phosphorea* and *A. murrayi* Luc proteins and the crystal structure of *R. reniformis* Luc complexes with coelenteramide (CEI) oxyluciferin (PDB ID: 7OMR). (E) Superimposed spectrum of the *in vivo* luminescence of *A. murrayi* (green) and the *in vitro* luminescent assay of *A. murrayi* luciferase in the presence of coelenterazine (fc.: 6 µM) (blue).

The retrieved sequence (CL1333; **Figure 4B** ) of *P. phosphorea* luciferase has an estimated molecular weight of 35.84 kDa. A comparison of the amino acid sequences of *P. phosphorea* and anthozoan luciferases demonstrated the presence of the catalytic triad involved in luciferase activity (**Figure S3; File S1** ). These key sites consist of an aspartate residue in position 120, a glutamate residue in position 144, and a histidine in position 285. Residues asparagine 53 and tryptophan 121 (N53 and W121), previously established for the O_2_ stabilization and the CO_2_ release within *Renilla* luciferase [14], were also retrieved in our luciferase sequences (**Figure S3**). The retrieved *P. phosphorea* luciferase sequence, based on RNA-seq data, appears to be highly similar to other known anthozoan luciferases. It shares 90.26% identity and 95% similarity with *Renilla reniformis* luciferase (*Renilla*-luciferin 2-monooxygenase). This sequence was validated by DNA amplification and sequencing using *R*Luc primers. FPKM analyses revealed that this sequence was mostly expressed in the pinnule and rachis (**Figure 4B**).

The *A. murrayi* luciferase (Comp24811, **Figure 4C**), with a molecular weight estimated at 34.61 kDa and consisting of 304 amino acids, exhibits 58% sequence identity and 98% coverage compared to *R*Luc, highlighting significant similarities. Sequence alignment of *A. murrayi* luciferase with *R*Luc has revealed the conservation of the catalytic triad and active site. The *R*Luc structure features two distinct domains: a cap domain and alpha/beta-hydrolase domain. Key residues essential for enzymatic activity, including the substrate entry tunnel and catalytic triad (D120, E144, and H285), are located in the cap domain (**Figure S3** ). These elements have also been identified in the luciferase sequence of *A. murrayi*, as reported by Rahnama et al., 2017 [62] and Khoshnevisan et al. 2018 [63] for *Renilla* luciferase. Through the expression of recombinant luciferase in *Escherichia coli* using degenerate primers derived from *A. murrayi* transcriptome analysis, *A. murrayi* luciferase was tested for preliminary downstream expression, yielding an active enzyme capable of producing blue light (λmax = 498 nm) upon coelenterazine addition (**Figure 4E**).

Structural models of *P*. *phosphorea* and *A*. *murrayi* LUCs show a canonical aba-sandwich fold with a helical cap domain (**Figure 4D**). Overall comparison between the crystal structure of *R*. *reniformis* LUC complexed with coelenteramide (CEI) oxyluciferin and structural models of *P*. *phosphorea* and *A*. *murrayi* LUCs showed root-mean-square deviation (RMSD) on the C_a_-atoms of 0.7 and 1.3, respectively. Careful inspection of the modeled LUC structures revealed that key residues of the catalytic pentad are conserved and properly positioned for productive catalysis (**Figure 4D** ). From these, three residues (aspartate-histidine-glutamate) function as a protein-relay system protonating a CEI oxyluciferin at an amide nitrogen, while the two residues (asparagine and tryptophan) are responsible for co-substrate (dioxygen) binding. Moreover, an aspartate residue, responsible for tuning the emission wavelength, found on the rim of the catalytic pocket in *R*. *reniformis* luciferase [14], is also conserved in these luminescent species.

Several sequences coding for GFP-like sequences were retrieved from *the P. phosphorea* and *A. murrayi* transcriptomes. *P. phosphorea* GFP (CL380; 20.81 kDa) and *A. murrayi* GFP (Comp24361; 27.24 kDa) appear homologous to other anthozoan sequences, and both sequences clustered with GFP sequences of bioluminescent Scleralcyonacea (**Figure 5A; Figure S3; File S2** ). *P. phosphorea* GFP shares 79.42% identity and 86% similarity with the GFP of *R. reniformis*. In comparison, *A. murrayi* GFP shared 73.06% identity and 87% similarity with the GFP of *Cavernularia obesa*. Pinnules and rachises were the main sites of *P. phosphorea* GFP expression (**Figure 5B** ). Interestingly, a perfect sequence repetition was observed in the most expressed *P. phosphorea* GFP sequence (**Figure S3** ). Among the unigene pairs found in the *A. murrayi* transcriptome, Comp24361 appeared to be more highly expressed (**Figure 5C**). Sequence analysis pinpointed a lack of N-terminal part for the *Anthoptilum* sequence (**Figure S3**).

**Figure 5.**
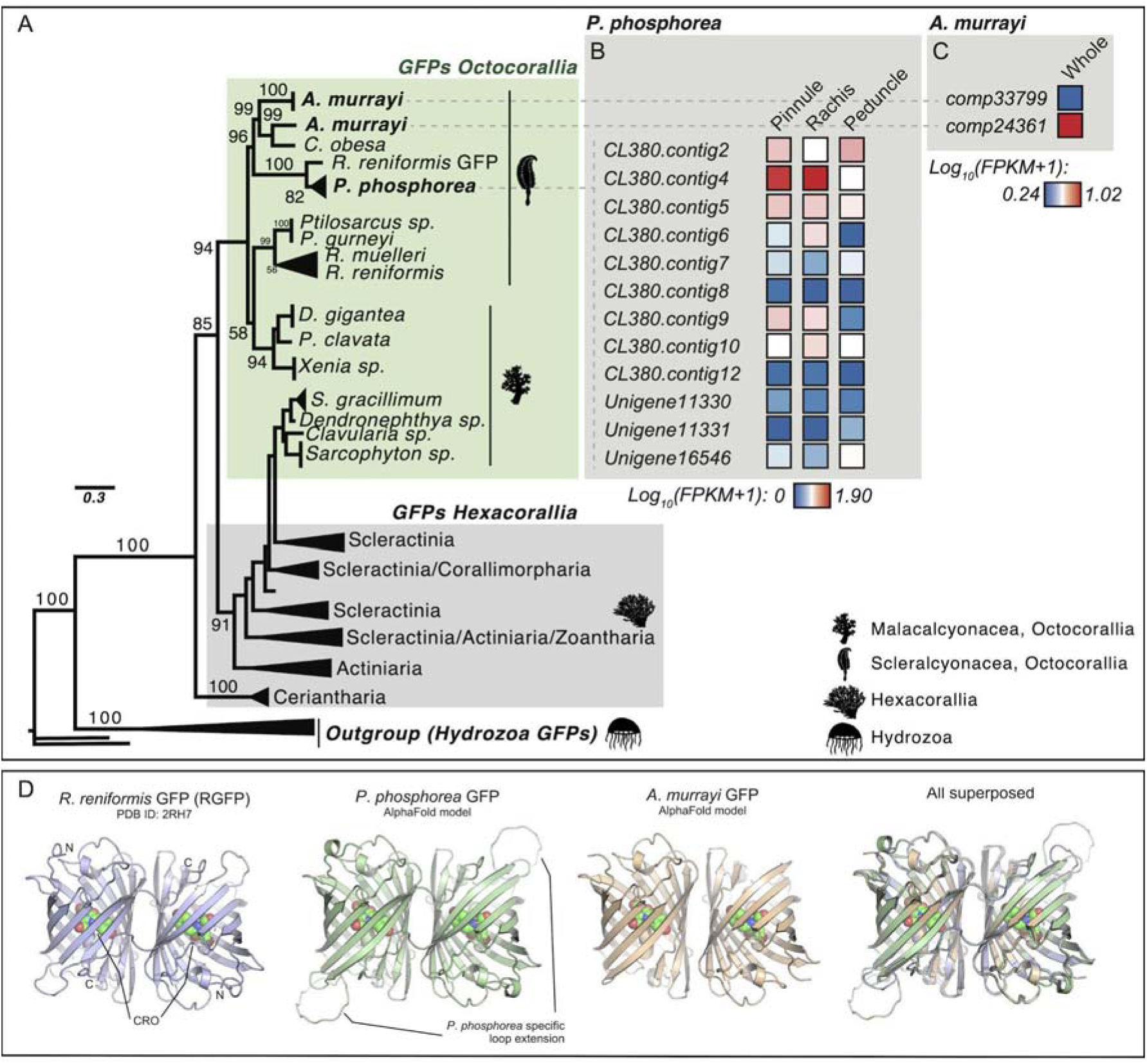
Phylogenetic tree of anthozoan fluorescent proteins, including *P. phosphorea* and *A. murrayi* amino acid sequences. (A) Maximum likelihood tree based on green fluorescent protein amino acid sequence alignment. The tree was calculated by the IQ-tree software using the WAG+R4 model of evolution. Numbers at the nodes indicate ultrafast bootstrap percentages based on 1,000 replicates. The scale bar represents the percentage of amino acid substitutions per site. Hydrozoan GFP sequences were used to root the tree. (B) The expression level of each retrieved GFP sequence for a distinct portion of *P. phosphorea*. (C) The expression level of each retrieved GFP sequence in *the whole A. murrayi* specimen. (D) Structural comparison of AlphaFold models of *P. phosphorea* and *A. murrayi* GFP proteins and the crystal structure of *R. reniformis* GFP (PDB ID: 2HR7).

Structural models of *P*. *phosphorea* and *A*. *murrayi* GFPs show a characteristic /3-barrel fold with a fluorophore moiety (CRO) covalently bound inside the barrel (**Figure 5D**). *In silico* modeling suggests that these proteins may associate as homodimers, similar to the homodimeric structure observed in the crystal structure of *R*. *reniformis* GFP. However, further experimental validation is required to confirm this hypothesis. Comparison between the crystal structure of *R*. *reniformis* GFP and models of *P*. *phosphorea* and *A*. *murrayi* GFPs showed RMSD on the C_a_-atoms of 0.9 and 1.1, respectively, highlighting their high similarities. A structural feature distinguishing *P*. *phosphorea* and *A*. *murrayi* GFPs from their *R*. *reniformis* counterpart is the composition of the fluorophore moiety. While *R*. *reniformis* fluorophore is generated from a serine-tyrosine-glycine tripeptide, the fluorophores of *P*. *phosphorea* and *A*. *murrayi* are formed from a glutamine-tyrosine-glycine tripeptide, which may affect the fluorescent properties of these proteins. Moreover, there is one markedly extended solvent-exposed loop in *P*. *phosphorea* GFP (**Figure 5D** ). We believe that this species-specific extension might affect protein–protein complexation during the radiationless resonance energy transfer process, but future experimental evidence is needed to verify this hypothesis.

Different sequences of CBPs and CBPs-like were retrieved from *the transcriptomes of P. phosphorea and A. murrayi*. Some of these sequences clustered with the CBPs of luminous Scleralcyonacea, whereas others were found to be clustered with photoprotein-like proteins (**Figure 6A; File S3** ). One sequence appeared to be mainly expressed within the *P. phosphorea* pinnule and rachis (CL1544; 26.84 kDa) and *the A. murrayi* colony (Comp 22336; 21.08 kDa) (**Figure 6B, C** ). The most highly expressed *P. phosphorea* CBP appeared highly similar to other luminous anthozoan sequences. *P. phosphorea* CBP shared 85.33% identity and 94% similarity with the luciferin-binding protein of *R. reniformis*, while *A. murrayi* CBP shared 50% identity and 74% similarity with the luciferin-binding of *R. reniformis*. Interestingly, sequences clustered in the photoprotein-like protein group appeared to be mainly expressed within the non-photogenic peduncle tissue in *P. phosphorea*.

**Figure 6.**
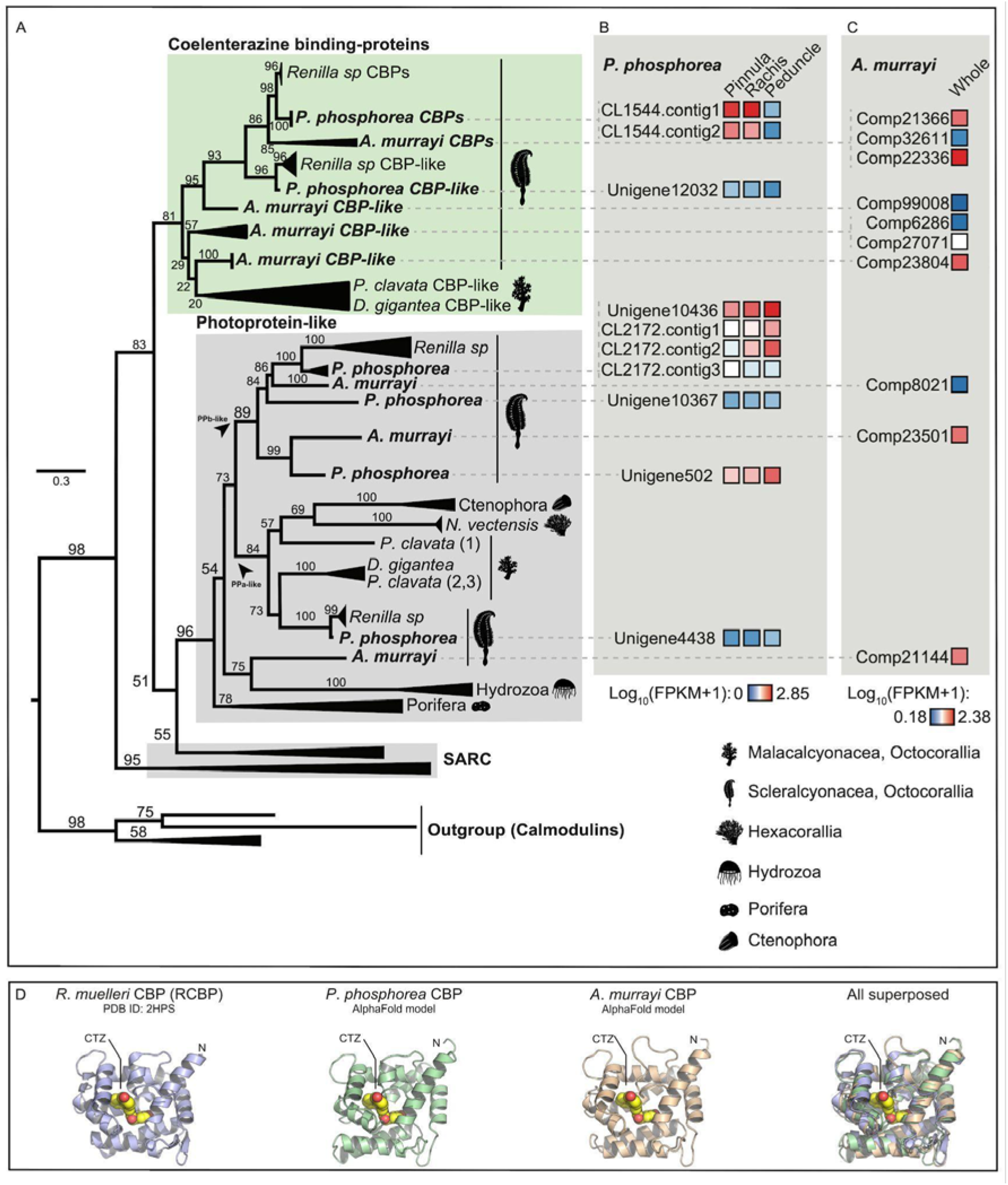
Phylogenetic tree of coelenterazine-binding proteins, including *P. phosphorea* and *A. murrayi* amino acid sequences. (A) Maximum likelihood tree based on the amino acid sequence alignment of coelenterazine-binding proteins. The tree was calculated by the IQ-tree software using the LG+R4 model of evolution. Numbers at the nodes indicate ultrafast bootstrap percentages based on 1000 replicates. The scale bar represents the percentage of amino acid substitutions per site. Calmodulin sequences were used to root the tree. (B) The expression level of each retrieved CBP and CBP-like sequence for a distinct portion of *P. phosphorea*. (C) Expression level of each retrieved CBP and CBP-like sequence in *the whole A. murrayi* specimen. (D) Structural comparison of AlphaFold models of *P. phosphorea* and *A. murrayi* CBP proteins and the crystal structure of *R. muelleri* CBP complexes with coelenterazine (CTZ) luciferin (PDB ID: 2HPS).

Finally, structural models of *P*. *phosphorea* and *A*. *murrayi* Ca^2+^-regulated CBPs reveal a typical EF-hand fold, containing three Ca^2+^-binding sites. As shown in **Figure 6D** , comparison between the crystal structure of *R*. *reniformis* CBP complexed with coelenterazine (CTZ) luciferin and structural models of *P*. *phosphorea* and *A*. *murrayi* LUCs show a structural similarity, with RMSD on the C_a_-atoms of 2.3 and 3.2, respectively. The RMSD values were higher than those observed for LUC and GFP proteins, but this is likely caused by a large conformational space that is searched by CBP proteins. Importantly, our modeling reveals an internal cavity in *P*. *phosphorea* and *A*. *murrayi* CBPs that is capable of accommodating CTZ luciferin, suggesting that these proteins are indeed functional luciferin-binding proteins (**Figure 6D**).

### Luciferase expression and green autofluorescence in Pennatula phosphorea and Funiculina quadrangularis

Based on the high sequence similarity of *P. phosphorea* luciferase with *Renilla* luciferase, a commercial anti-Renilla luciferase antibody was selected for immunodetection. Immunoblot analyses revealed strong anti-luciferase immunoreactive bands in both extracts of the pinnule and rachis tissues (**Figure S4** ). The bands correspond to a protein with an approximate molecular weight of 35 kDa, matching the molecular weight of the predicted *P. phosphorea* luciferase and the *Renilla* luciferase molecular weight of 36 kDa. No labeling was detected in the peduncle tissue extract (**Figure S4**).

On autozooid polyps of the pinnules (**Figure 7A-D** ), a strong green autofluorescent signal was observed at the tentacle crown base before (**Figure 7B** ) and after (**Figure 7C** ) paraformaldehyde fixation. This fluorescent signal is located in clusters of cells at the tentacle junctions. Strong anti-*Renilla* luciferase immunoreactivity was observed at the same level as the green fluorescence signal (**Figure 7C, D** ). On siphonozooid polyps of the rachis, the autofluorescent signal was also observable and was located as green dots (from 10 to 25 µm diameter) spread in the tissue, generally in pairs (**Figure 7E** ). Anti-*Renilla* luciferase-positive cells colocalized with these autofluorescent dots (**Figure 7E, F** ). Finally, no green fluorescence or immunolabeling was detected in peduncle tissue (data not shown).

**Figure 7.**
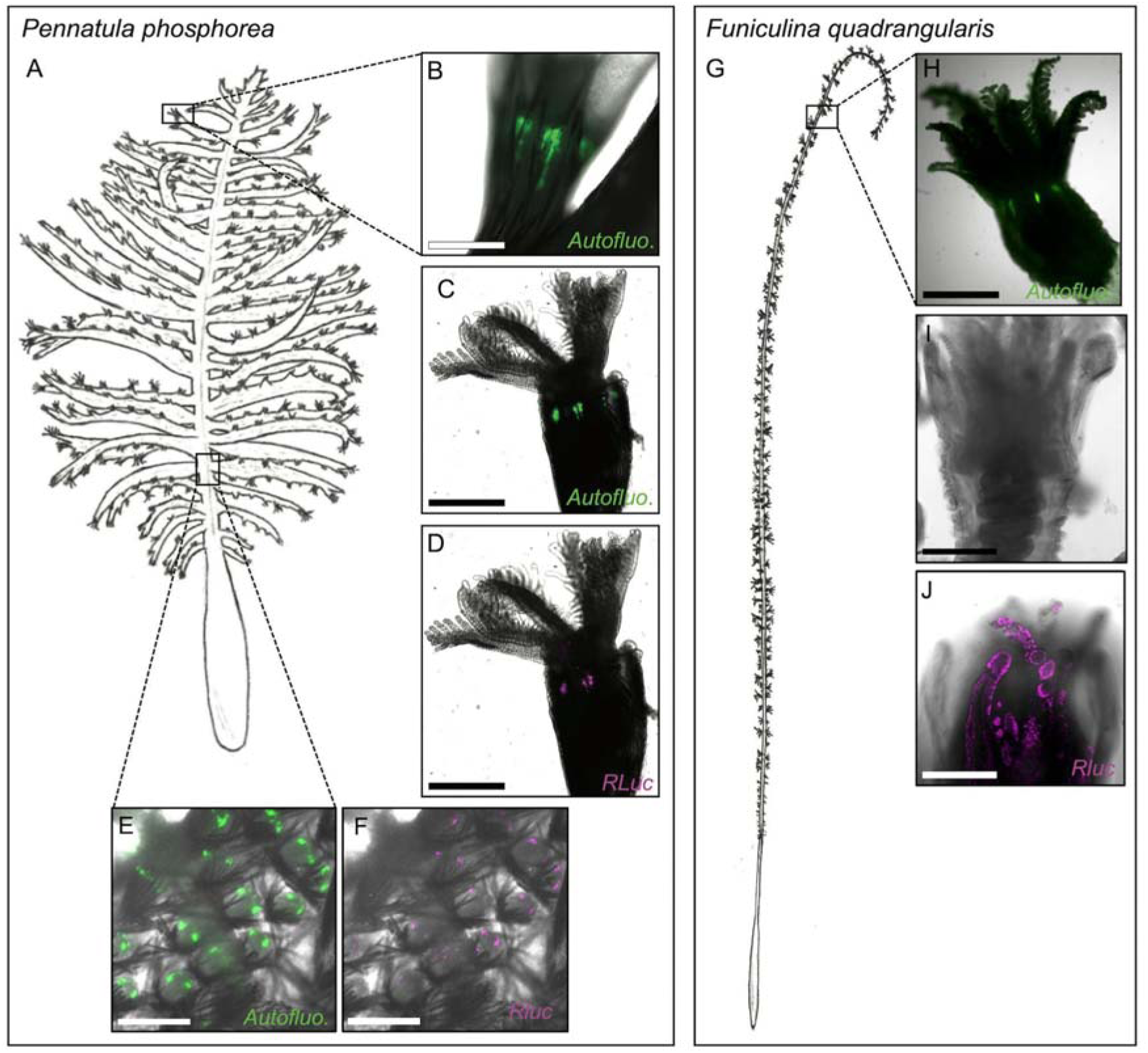
Autofluorescence and luciferase immunodetection in *P. phosphorea* and *F. quadrangularis.* (A) Schematic illustration of *P. phosphorea*. Natural green autofluorescence (B), green fluorescence after fixation (C), and luciferase immunodetection (D; magenta) of the *P. phosphorea* pinnule autozooids. Green fluorescence after fixation (E) and luciferase immunodetection (F; magenta) in *P. phosphorea* rachis siphonozooids. (G) Schematic illustration of *F. quadrangularis*. Natural green autofluorescence (H), observation after fixation with no autofluorescent signal (I), and luciferase immunodetection (J; magenta) of *the F. quadrangularis* autozooids. Scales: B-F, H - 500 μm; I, J - 250 μm.

For *Funiculina* polyps (**Figure 7G**), a green autofluorescent signal was observed in the freshly dissected specimens before fixation (**Figure 7H** ). This green autofluorescence completely disappeared after fixation (**Figure 7I**). Luciferase localization differed from that in *P. phosphorea*. A strong luciferase signal was detected within the polyp tentacle tissue and not at the base of the polyp crown (**Figure 7I, J**).

Negative controls with the omission of the primary antibody did not reveal any non-specific binding of the secondary antibodies (data not shown).

### Calcium is involved in the bioluminescence of Pennatula phosphorea and Funiculina quadrangularis

Based on *P. phosphorea* coelenterazine-binding protein retrieval in the *Pennatula* transcriptome data and the literature mentioning the potential implication of Ca^2+^ in the release of luciferin from coelenterazine-binding proteins, Ca^2+^ involvement in the light emission process of *P. phosphorea* and *F. quadrangularis* was investigated. The tested specimens of both species revealed a drastic increase in light production when immersed in ASW with a doubled Ca^2+^ ion concentration (20 mM) (**Figure 8A, B** ). At the same time, no statistical differences were observed between the normal ASW (10 mM) and ASW devoted to Ca^2+^ ions (0 mM) (**Figure 8A, B**). Analysis of the effects of the A23187 ionophore showed no statistical differences compared to the calcium concentration (**Figure 8C, D** ). For both specimens, adrenaline triggered light production when calcium was present in ASW (**Figure 8E, F**). While an increase in the mean Ltot was observed between adrenaline tested samples at 10 mM and 20 mM of CaCl_2_ in the medium, this increase was not statistically supported (**Figure 8E, F**). Finally, no significant differences were observed in Ltot after KCl application at the three Ca^2+^ concentrations (**Figure S5**).

**Figure 8.**
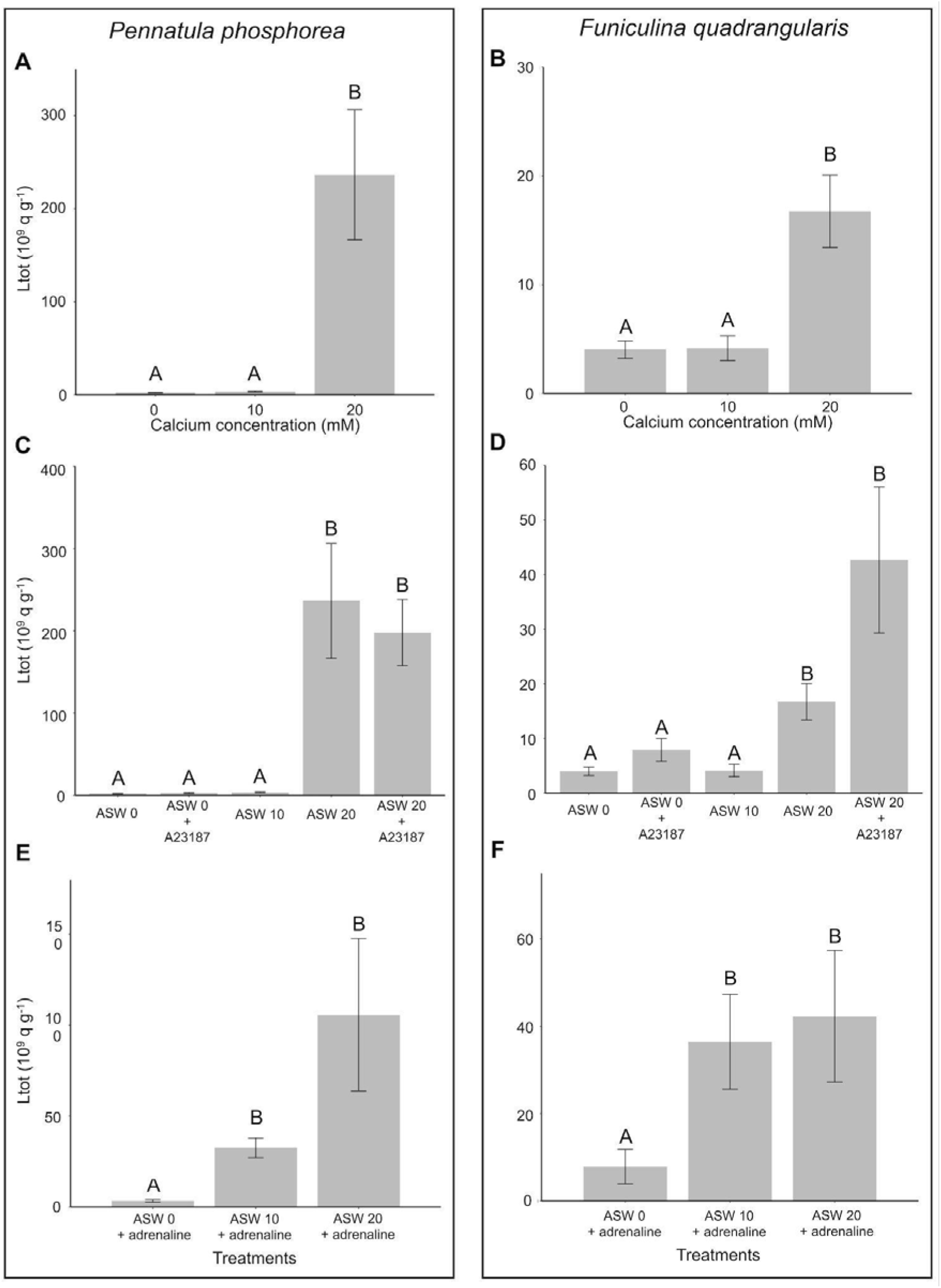
Calcium involvement in *P. phosphorea* and *F. quadrangularis* light emissions. Experiments were performed on *P. phosphorea* (A, C, E) and *F. quadrangularis* (B, D, F). Effect of different concentrations of calcium (0, 10, 20 mM) in the medium on the total light emission (Ltot) (A, B). Effect of different calcium concentrations (0, 10, 20 mM) in the medium in the presence and absence of the calcium ionophore A23187 on the Ltot (C, D). Effect of different concentrations of calcium (0, 10, 20 mM) in the medium in the presence of adrenaline (10 mol L ) on the Ltot (E, F). different lettering indicates statistical differences.

## Discussion

Through biochemical cross-reaction experiments, this study demonstrated that the bioluminescence of *P. phosphorea, F. quadrangularis*, and *A. murrayi* is associated with a coelenterazine-dependent luciferase homologous to the well-known luciferase of the sea pansy *Renilla*. The measured concentration of coelenterazine in *P. phosphorea* pinnules closely aligns with the reported values of 141 and 282.4 ng g^-1^ for the phylogenetically closely related species *R. muelleri* and *C. obesa*, respectively [1]. Similarly, the mean luciferase activity of *P. phosphorea* pinnules reaches a comparable range as other organisms using *Renilla*-type luciferases such as *R. muelleri* (± 130 10^9^ q g^-1^ s^-1^) and *C. obesa* (± 220 10^9^ q g^-1^ s^-1^), or the sympatric ophiuroid *Amphiura filiformis* (± 69 10^9^ q g^-1^ s^-1^) [1,64]. While the luciferase activity recorded for *F. quadrangularis* is higher than *P. phosphorea*, this species exhibits a smaller coelenterazine content, being closer to recorded values of the echinoderms *A. filiformis* (5.4 ng g^-1^; [64]) or the crinoid *Thalassometra gracilis* (Carpenter, 1888) (4.5 ng g^-1^; [65]). Transcriptome and phylogenetic analyses confirm results for *P. phosphorea* and *A. murrayi* with a clear sequence conservation of the retrieved luciferases with the *Renilla* luciferase. Consistent with the literature, species such as *D. gracile* or undetermined species from the genera *Umbellula, Pennatula*, and *Funiculina* also present evidence of the use of coelenterazine as substrate and a *Renilla*-like luciferase as enzyme of their bioluminescent systems [13].

Our phylogenetic analysis underlines three distinctive clades of *Renilla*-like luciferase. One of these clades (Clade A) contains all the known sequences of anthozoan light-emitting luciferases. The other two clades (Clades B and C) contain other homologous *R*luc-like sequences, notably from non-luminous species. The related biochemical functionality and bioactivity of these B and C clades sequences are unknown and need further investigations to fully apprehend the evolution of luciferase among anthozoans. Nevertheless, this clustering led to hypothesized duplications of the ancestral gene with either (*i*) a neo functionality as a “real” functional luciferase able to catalyze a bioluminescent reaction or (*ii*) a bi-functionality of these enzymes as light-producing enzyme and another ancestrally conserved enzymatic function. The ancestral functionality of the *R*Luc enzyme has been assumed to originate from bacterial haloalkane dehalogenases, enzymes with a hydrolase activity cleaving bonds in halogenated compounds [66]. A basal gene transfer from bacteria until metazoans has been hypothesized [18,66]. Interestingly, *R*Luc-like luciferases have been demonstrated to be the enzymes involved in the bioluminescence of phylogenetically distinct species such as the brittle star *A. filiformis* [18] and the tunicate *P. atlanticum* (Péron, 1804) [19], letting assumed that the sequence was convergently coopted multiple times during the evolution. Consistent with previous research on *R*Luc sequences, our retrieved *P. phosphorea* and *A. murrayi* luciferase (Clade A) present high sequence similarity with the other pennatulaceans functional luciferases, also revealing conservation of the catalytic triads essential for the luciferase catalytic activity [14,51,67]. The complete characterization of *P. phosphorea, A. murrayi*, and *F. quadrangularis* luciferases will allow us to better apprehend the functionality and evolution of these luciferase enzymes. Future research will enable us to better characterise these luciferases and also the interactions between these enzymes and their substrates and associated molecules, as is the case with the approach to the role of *Renilla* luciferase residues (e.g., mutagenesis, QM/MM [14,51,67]).

While the retrieved GFP sequences are unique and well clustered with other Scleralcyonacea sequences, a similar observation, as for the luciferases, occurs for the CBPs and CBPs-like with multiple retrieved sequences clustered in different groups. The first group represents the functional coelenterazine-binding protein retrieved and characterized in luminous species, while the second group, named photoprotein-like proteins, raise questions on the exact functionality of these retrieved sequences. These photoprotein-like proteins might also be involved in calcium binding. Active photoproteins of cnidarians and ctenophores depend on calcium to trigger light emission [68–71]. Nevertheless, the higher expression within the *P. phosphorea* peduncle assumed another function without a relationship with the bioluminescence.

AlphaFold is a neural network machine learning tool for predicting macromolecular structures and complexes, providing structural models with near-atomic accuracy even in the absence of known similar structures [72,73]. Here, we employed AlphaFold to predict macromolecular structures of the most expressed genes encoding for LUC, GFP, and CBP proteins in *P*. *phosphorea* and *A*. *murrayi*. Computational predictions yielded structural models with very high confidence scores (>90) for all analyzed proteins and that are structurally similar to crystallographic structures of well-characterized LUC [14], GFP [12], and CBP [74] proteins encoded by the sea pansies *R*. *reniformis* and *R*. *muelleri*. Taken together, our structural predictions suggest that the identified *P*. *phosphorea* and *A*. *murrayi* LUC, GFP, and CBP proteins with the highest expression values are structurally and functionally relevant and responsible for bioluminescence in these species.

Kept in captivity without exogenous coelenterazine supply, *P. phosphorea* can still produce light after one year, even if the coelenterazine content and luciferase activity decrease. Therefore, these results support a *de novo* synthesis of the coelenterazine substrate in *P. phosphorea*. Coelenterazine *de novo* synthesis by luminous marine organisms has been documented for the calanoid copepods *Metridia longa* (Lubbock, 1854) and *M. pacifica* (Brodsky, 1950), the oplophorid shrimp, *Systellaspis debilis* (Milne-Edwards, 1881), and two ctenophores, *Mnemiopsis leidyi* (Agassiz, 1865) and *Bolinopsis infundibulum* (Müller, 1776) [75–78]. The natural precursors of coelenterazine have been demonstrated to be the L-tyrosine and L-phenylalanine amino acids in *M. pacifica* [77]. By screening the transcriptomic data of 24 ctenophores, Francis et al., 2015, assumed the involvement of a non-heme iron oxidase-like enzyme, similar to isopenicillin-N-synthase, in the biosynthesis pathway of this luciferin [79]. Future research could be carried out to validate the *de novo* biosynthesis of the coelenterazine bioluminescent substrate, in particular by maintaining *P. phosphorea* and their offspring over generations in captivity in the same conditions without coelenterazine supply [78,80]. This protocol could be applied to other pennatulaceans to determine whether the coelenterazine genesis is a common trait in this clade. Nevertheless, the first challenge would be to control the life cycle of these species in captivity. Similarly, it would be interesting to analyze different pennatulacean transcriptomes searching for enzymes homologous to the isopenicillin-N-synthase potentially involved in the coelenterazine biosynthetic pathway, as retrieved in ctenophores.

In *P. phosphorea*, the morphological localization of luciferase expression matched the green fluorescent sites obtained in unfixed and fixed tissues. Green autofluorescence observed on unfixed specimens is assumed to be a mix of the native autofluorescence of coelenterazine and the autofluorescent reaction occurring through the GFP, while green autofluorescence observed after tissue fixation corresponds only to the GFP signals. Comparatively, the *in vivo* green fluorescence observed in the *F. quadrangularis* tissues, which disappeared after fixation, was assumed to be related only to the natural autofluorescence of coelenterazine and the lack of GFP in this species. In contrast to *P. phosphorea,* which emits green waves of light at λmax = 510 nm, *F. quadrangularis* emits blue at λmax = 485 nm, supporting the absence of GFP for this species [13,36,61,81]. This natural autofluorescence has recently been demonstrated to appear and disappear from the photogenic site, depending on the substrate dietary acquisition of the brittle star *A. filiformis*. This species depends on the trophic acquisition of the coelenterazine substrate to produce light [64,82]. When the brittle star was fed with coelenterazine-containing food, green autofluorescent spots appeared at the level of spine-associated photocytes [82]. As for *F. quadrangularis*, a similar disappearance of the green fluorescent signal (possibly attributed to the coelenterazine) has been observed in the fixed tissue of the brittle star species [82]. The autofluorescent sites observed along the tentacle bases of the autozooids of *P. phosphorea* are consistent with the already described location of the autofluorescent photogenic cell processes in autozooids of *Stylatula elongata* [83], and to a lesser extent, *Acanthoptilum gracile,* and *Renilla koellikeri* [83,33]. For the former species, it was noticed that the photocytes process followed the same orientation as the longitudinal muscles, allowing autozooids to retract [33]. On the other hand, the luciferase expression site in *F. quadrangularis*, in the cellular processes of the apical part of the tentacle, was never reported before. This location along the polyp tentacles matches the described position of photocytes in autozooids of *Ptilosarcus* species [83].

Our spectrum measurement performed with the *A. murrayi* recombinant luciferase is consistent with already observed coelenterazine-dependent systems in other species emitting in the same range of wavelengths. Recorded spectrum for coelenterazine-dependent luciferase systems can vary between 475 to 493 nm in *P. atlanticum*, 482 nm emitted by *R*Luc, 472 nm emitted by the brittle star *A. filiformis*, 455 nm emitted by *Oplophorus gracilirostris* and 485 nm emitted by *F. quadrangularis* [18,19,66,84]. According to the natural spectrum recorded on the whole specimens (λmax = 513 nm), similar to the spectrum measured for *P. phosphorea* [61], and the transcriptomic presence of a GFP sequence in *A. murrayi*, this species is strongly assumed to display a GFP-associated coelenterazine-dependent luminous system. At least six anthozoan species present expression of a GFP and the production of green light. Some of these species may live in sympatry with anthozoan blue emitters, such as *P. phosphorea* and *F. quadrangularis,* with an overlap of depth repartition and habitat preferences occurring. Therefore, questions arise concerning luminescence’s exact function(s) among anthozoans. Even if some assumptions are proposed in the literature, no one has ever developed an ethological protocol to validate them [37]. Therefore, why some anthozoan evolved green light emission while the primary coelenterazine-luciferase reaction produces blue light remains. Different function(s) in the same environment may have led to the acquisition or the loss of the GFP gene by some anthozoan species upon evolutionary constraints.

As demonstrated for other pennatulacean species, CBP seems to be an essential component of the luminous system [21–25,27]. The retrieved CBPs in *P. phosphorea* and *A. murrayi* are congruent with this literature. Gene ontology distribution analyses performed on the different tissues of *P. phosphorea* underline a high expression of calcium ion binding proteins, including the retrieved CL1333 CBP, in the photogenic tissue (pinnule and rachis) of this species. The expression of this gene within the photogenic tissues supports its involvement in the luminous reaction. As demonstrated for *Renilla* by Stepanyuk et al., 2008, CBPs need calcium as a cofactor to release the coelenterazine [29]. In addition to the retrieved CBPs sequence in the *Pennatula* transcriptome, our calcium assay results highlight the involvement of the calcium ion in the light emission process of both *P*. *phosphorea* and *F. quadrangularis*. Results obtained for the calcium ionophore A23187 reveal that the ion action does not result from intracellular calcium storage but rather is provided by external calcium input. Moreover, calcium is shown to be essential for the physiological luminescent response through adrenaline application. Pieces of evidence of calcium involvement in anthozoan luminescence were already described for *R. reniformis* and *V. cynomorium* [27,28].

These results let us assume a conservation of the coelenterazine-dependent *Renilla*-like luciferase bioluminescent system, involving also a CBP, across luminous Pennatulaceans (**Table S1**; [8]). The involvement of GFP appears species-dependent, with only a few species emitting blue light (**Table S1** ; [8]). Nevertheless, deeper investigations are needed to fully apprehend the conservation of those actors across the diversity of luminous pennatulaceans.

A hypothetical scheme of the generalized pathway could be established using our results and the literature on the pennatulacean luminescence mechanism (**Figure 9** ). The first step is the activation of catecholaminergic receptors through the binding of biogenic amines (mainly adrenaline and noradrenaline [35,36]), which will release an intracellular-associated G-protein [85,86]. G-protein could be involved in a large variety of intracellular pathways [87], some of which involve increasing intracellular calcium (via direct or indirect activation of calcium channels) [88–91]. This intracellular calcium increase will lead to the release of the coelenterazine through the binding of this ion on the CBP, leaving this luciferin free to react with luciferase in the presence of oxygen to produce blue light around 480 nm [9–11,16,21–25,27]. For pennatulaceans lacking GFP (*e.g., F. quadrangularis* ), this scheme ends here with the direct emission in blue color, while for those displaying GFP expression, the blue light is captured by this specific fluorescent protein and reemitted in green wavelength [22], such as for *P. phosphorea*. Future research is needed to validate this hypothetical scheme, and further investigations will be conducted to establish the functional activities of all these components in less-studied sea pens. Astonishingly, even if attempts were made on other anthozoans over the past decades, the *Renilla* bioluminescence system remains the only isolated and cloned system [14,16,22–25,27]. Despite the demonstrated widespread uses of the sea pansy bioluminescent system in biotechnology and biomedicine [*e.g.,* 92-94], the *Renilla* luciferase stands as the only one of the most commercially employed gene reporters in biomolecular sciences. Nonetheless, in the *Renilla* bioluminescence system, the exact action mode of CBP and calcium is not fully apprehended [29,95]. Considering these facts, our introspection into the bioluminescent system of other luminous pennatulaceans could be of great use for new biotechnological advances. Our results provided a better understanding of the evolution of the bioluminescence system and associated molecules from these enigmatic benthic sessile organisms.

**Figure 9.**
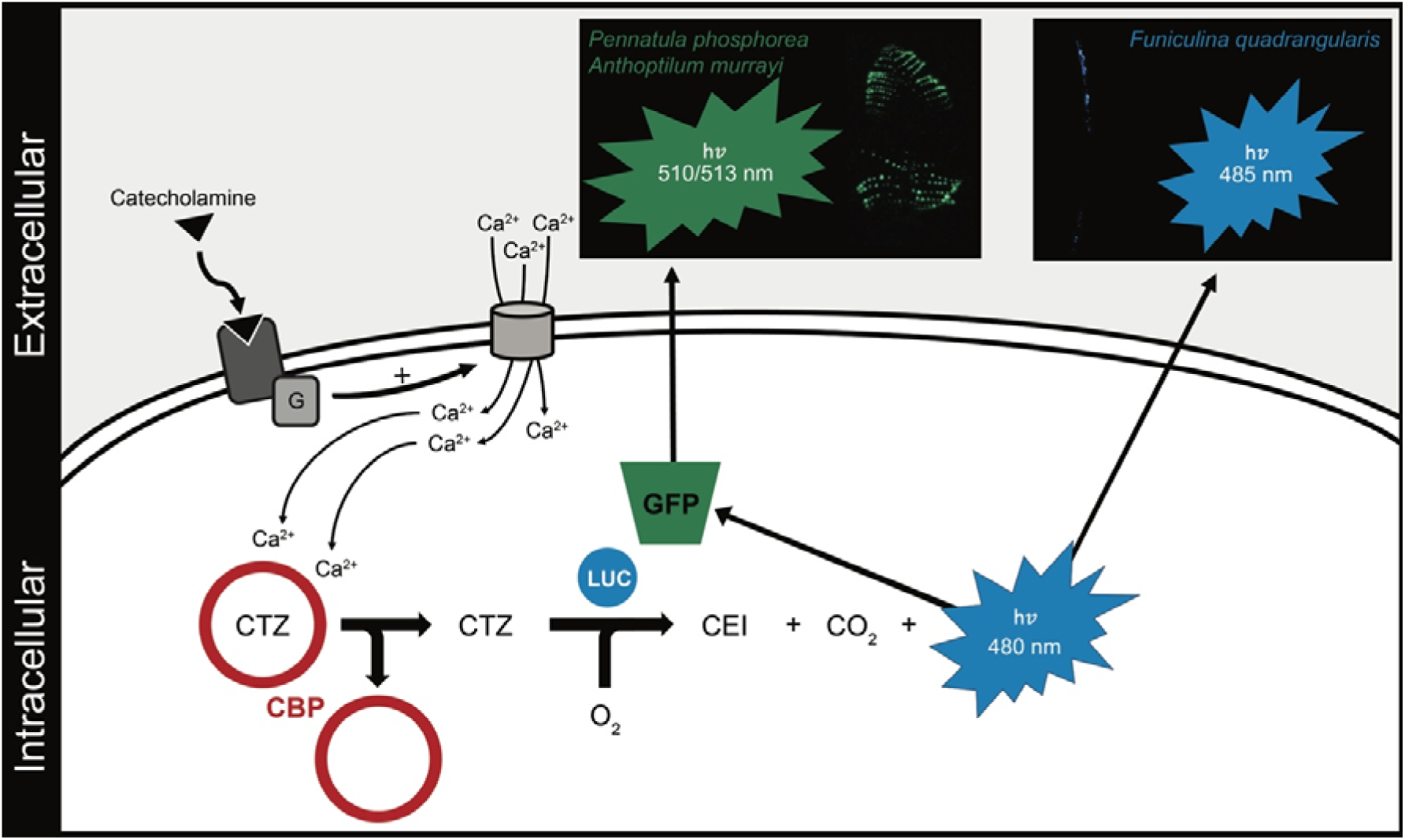
Schematic representation of the putative pathway driving the luminescence production in sea pens. Elements of this pathway have been compiled from the present results and the literature [13,14,17,21–25,32–36]. CBP, coelenterazine-binding protein; CEI, coelenteramide; CTZ, coelenterazine; G, G-protein; GFP, green fluorescent protein; LUC, luciferase. The clear colocalization of specific molecular actors (*e.g.,* catecholaminergic receptors, CBP, GFP, LUC) still needs to be established to confirm the following predicted pathway.

## Supplemental data captions

**Table S1. General overview of the bioluminescent knowledge among luminous anthozoans.** All data were extracted from the literature (only scientific data described for species down to the species level have been taken into account to minimize generalizations). CBP, coelenterazine-binding protein; GFP, green fluorescent protein.

**Table S2. Primers designed for *P. phosphorea* luciferase validation.**

**Table S3. Experiment protocol for the calcium assays.**

**Table S4. Description of the output sequenced data.** Q20 percentage is the proportion of nucleotides with a quality value larger than 20 in reads. GC percentage is the proportion of guanidine and cytosine nucleotides among total nucleotides.

**Table S5. Summary statistics of assemblies for *Pennatula phosphorea* pinnules, rachis, peduncle, and *Anthoptilum murrayi* transcriptomes.**

**Table S6. Transcript expression values (FPKM values) and public database sequences used during transcriptomic and phylogenetic analyses.**

**Figure S1: Luminometric measurements performed on the different area of *Pennatula phosphorea*.** (A) schematic representation of *P. phosphorea* with the different areas (upper, middle, lower) of the pinnules and rachis. (B) Mean coelenterazine content recorded and (C) luciferase activity for the different pinnule areas. (D) Mean coelenterazine content recorded and (E) luciferase activity for the different rachis areas.

**Figure S2: Biochemical assays following luminous parameters activity on *Pennatula phosphorea* pinnules (A, B, C) and rachis (D, E, F) during 1 year without coelenterazine supply.** Mean total light emission after KCl applications (A, D), mean coelenterazine content (B, E), and mean maximal light emission during luciferase activity experiments (C, D). Different lettering indicates statistical differences. The timing corresponds to experiments performed on wild-caught specimens (T0) after 6 months (T6) and twelve months of captivity (T12).

**Figure S3: Sequence alignments.** (A) Alignment of the retrieved luciferases with a highlight on the conserved catalytic sites. (B) Alignment of the retrieved green fluorescent protein sequences. (C) Alignement of the retrieved coelenterazine-binding protein sequences.

**Figure S4. Luciferase immunoblots on *Pennatula phosphorea* tissues (rachis, Ra; peduncle, Ped; and pinnule, Pin).**

**Figure S5. Total light emission (Ltot) after KCl applications with three different calcium concentrations in the medium.** Experiments were performed on (A) pinnules, and (B) rachis of *Pennatula phosphorea*. ASW, artificial seawater; 0, 10, 20 correspond to the calcium concentration (mM) in the medium.

**File S1**: FASTA file of the retrieved Clade A luciferase sequences

**File S2**: FASTA file of the retrieved green fluorescent protein sequences

**File S3**: FASTA file of the retrieved coelenterazine-binding protein sequences

## Acknowledgment

The authors acknowledge U. Schwarz, captain of the Alice vessel, and the skillful members of the Kristineberg Center (Goteborg University, Sweden) for their help during the *Pennatula* and *Funiculina* collection; and commandant J. Rezende and the crew of the R/V Alpha Crucis (Instituto Oceanográfico, USP). The authors also thank M. Jacquet for contributing to the study during his master’s thesis and C. Pels, ELIV laboratory technician who maintained the organisms in the aquaria at the Marine Biology Laboratory - UCLouvain. The authors also want to thank T. Wiegand from the mobile lab (TREC-EMBL) for her help in visualizing coelenterazine autofluorescence and immunolabeling in the wild-caught specimens. The authors also thank J. Mallefet for his helpful advice and help during the first organisms sampling and all along the experiments.

LD is a postdoctoral researcher at the Université de Louvain - UCLouvain, GG is a Ph.D. student at Universidade de São Paulo - USP, CC is a Ph.D. student under an FRIA fellowship, LB is a postdoctoral researcher at the Université de Louvain - UCLouvain, RR is an academic professor at UCLouvain, MRSM is an academic professor at Universidade de São Paulo - USP, MM is group leader at the Masaryk University, DTA is an adjunct professor at Federal University of ABC, SD is a Senior Lecturer and Associate professor at the University of Gothenburg, AGO is an assistant professor at Yeshiva University, and JD is a postdoctoral researcher at FNRS. This study is the contribution of BRC#422 of the Biodiversity Research Center (UCLouvain) from the Earth and Life Institute Biodiversity (ELIV) and the “Centre Interuniversitaire de Biologie Marine” (CIBIM).

## Data availability

Transcriptome raw reads were uploaded as Sequence Reads Archives (SRA): *A. murrayi* (PRJNA1144931), *P. phosphorea* (PRJNA1152785). The unigene annotation tables are accessible from the corresponding author upon request.

## Author contributions

LD, CC, and SD collected the *P. phosphorea* and *F. quadrangularis* samples, and GG and MRSM sampled *A. murrayi*. Data collection and analyses were performed by LD, CC, LB, GG, AO and JD. Transcriptome analyses were performed by LD, GG, DTA, AO, and JD. Phylogenetic analyses were performed by JD. Structural analyses were performed by MM. The project was supervised by RR, SD, AO, and JD. The original manuscript was written by LD, GG, CC, AO, and JD. All the authors reviewed and approved the final version.

