## Supplementary material for "Insights into the bioluminescence systems of three sea pens (Cnidaria: Anthozoa): from *de novo* transcriptome analyses to biochemical assays": Figure S1

A

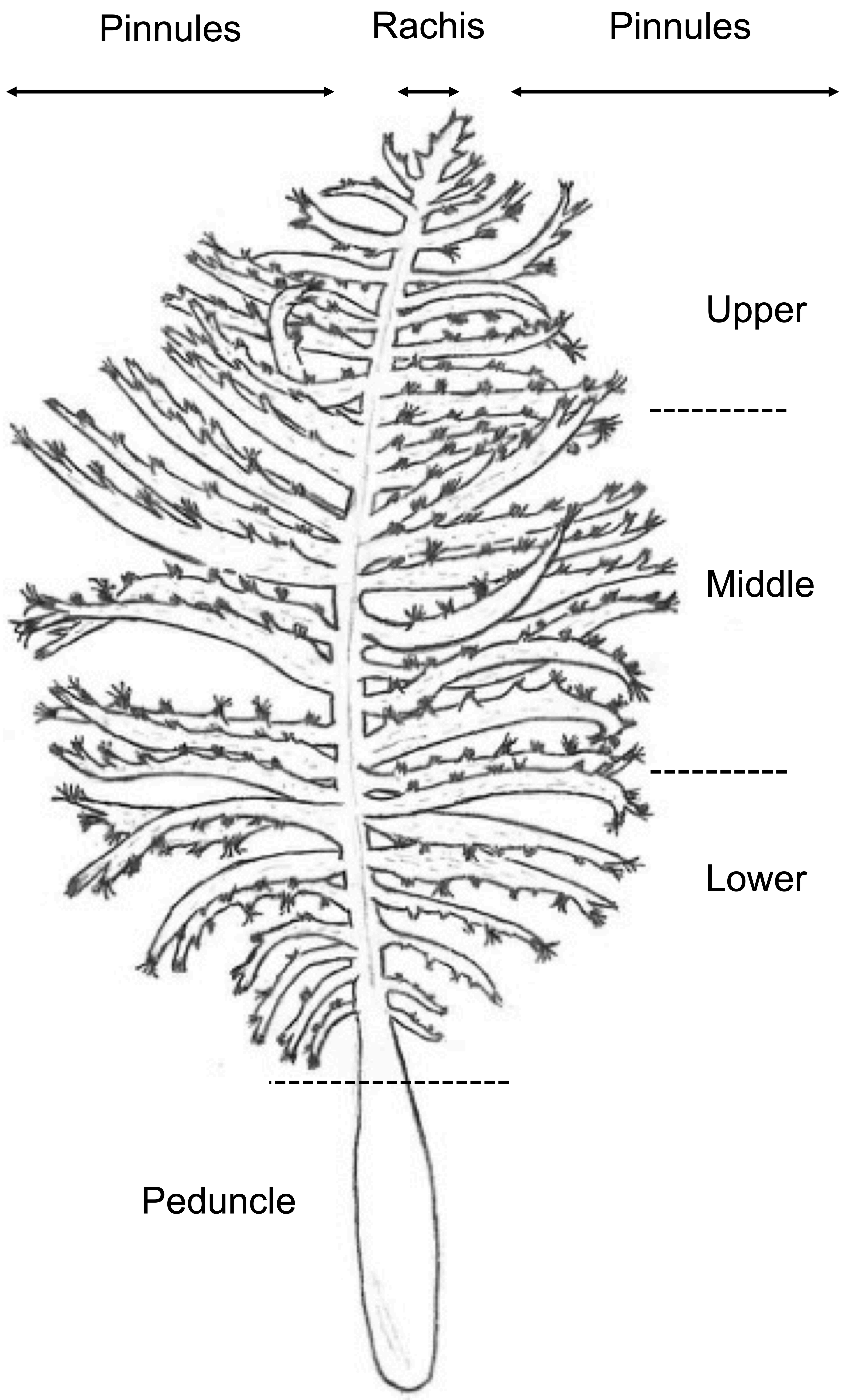


B

| Pinnule area | n | Mean coelenterazine content (ng g$^{-1}$) | SEM |
|---|---|---|---|
| Upper | 12 | 139.98 | 28.06 |
| Middle | 12 | 95.59 | 22.82 |
| Lower | 12 | 156.87 | 36.30 |

C

| Pinnule area | n | Mean luciferase activity ($10^9$ q g$^{-1}$ s$^{-1}$) | SEM |
|---|---|---|---|
| Upper | 12 | 152.3 | 28.6 |
| Middle | 12 | 148.1 | 27.8 |
| Lower | 12 | 88.0 | 9.9 |

D

| Rachis area | n | Mean coelenterazine content (ng g$^{-1}$) | SEM |
|---|---|---|---|
| Upper | 6 | 52.95 | 13.84 |
| Middle | 6 | 92.77 | 15.97 |
| Lower | 6 | 64.25 | 54.21 |

E

| Rachis area | n | Mean luciferase activity ($10^9$ q g$^{-1}$ s$^{-1}$) | SEM |
|---|---|---|---|
| Upper | 6 | 119.3 | 16.8 |
| Middle | 6 | 61.4 | 27.4 |
| Lower | 6 | 83.0 | 39.0 |
