## Supplementary figures and images for "Insights into the bioluminescence systems of three sea pens (Cnidaria: Anthozoa): from *de novo* transcriptome analyses to biochemical assays"

### Figure S2

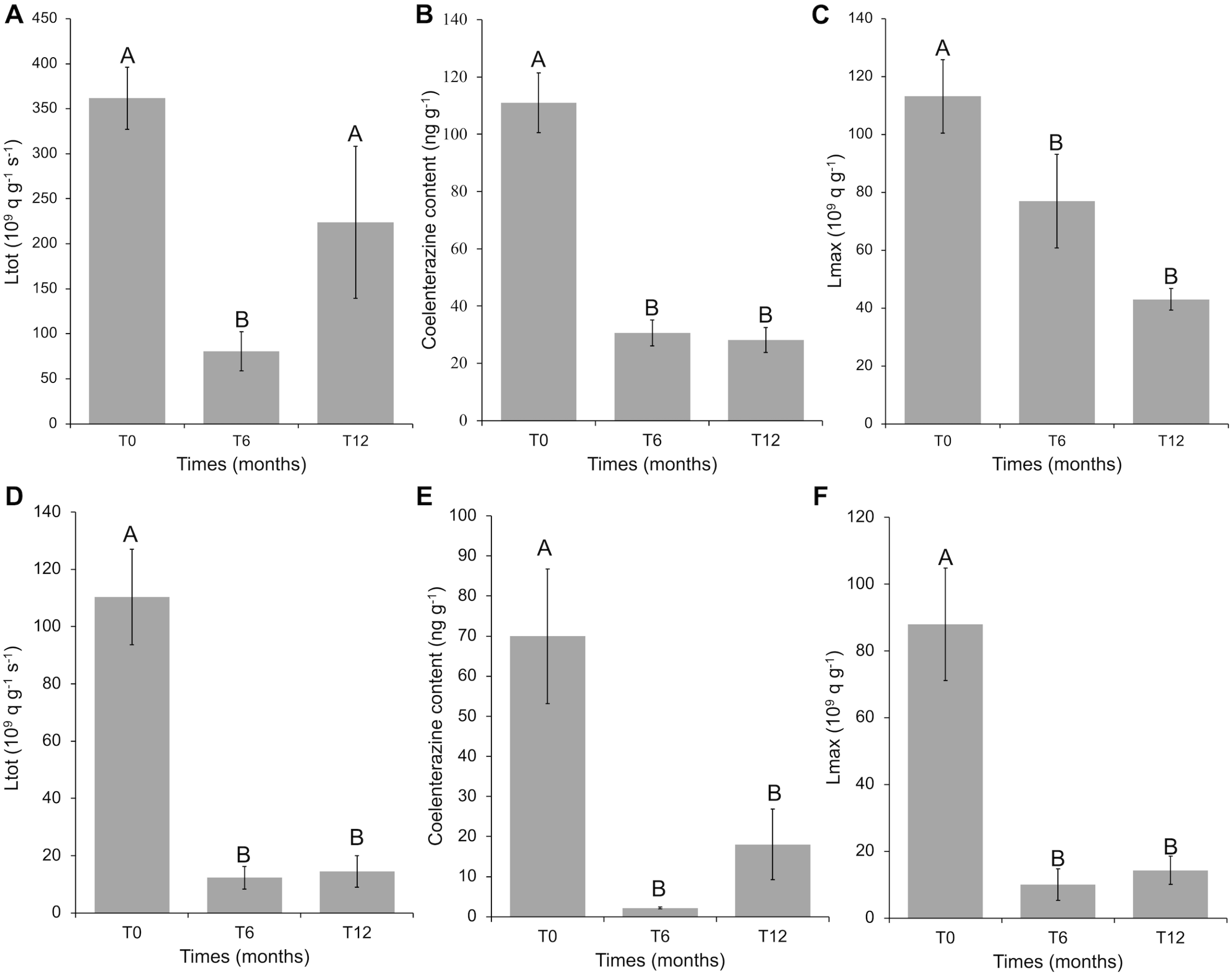

### Figure S3

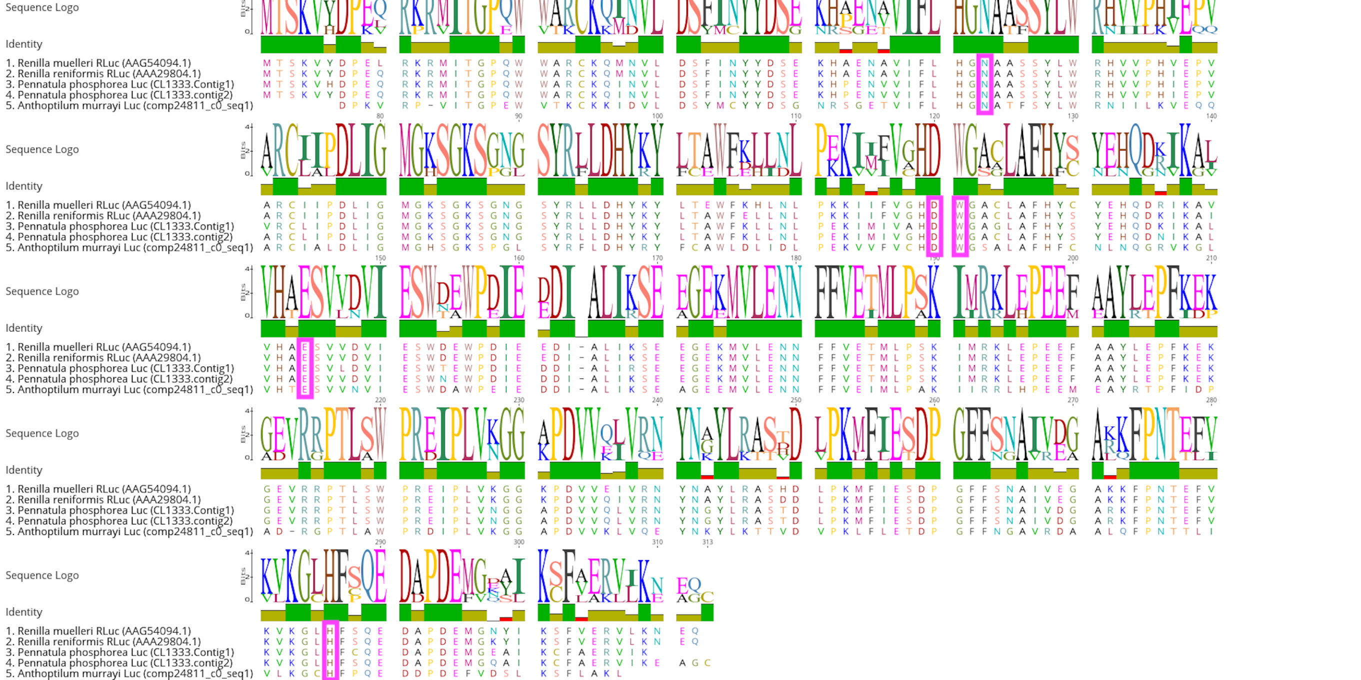

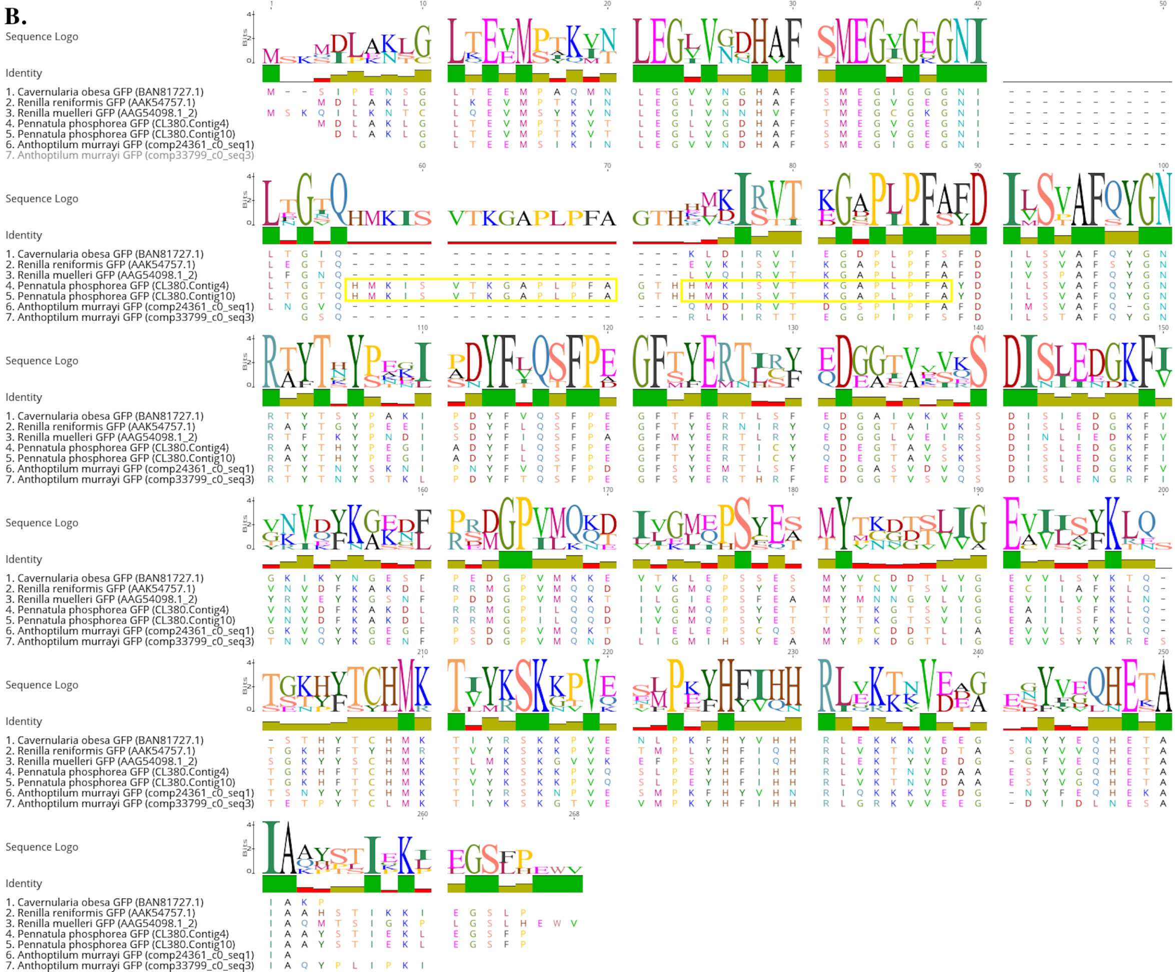

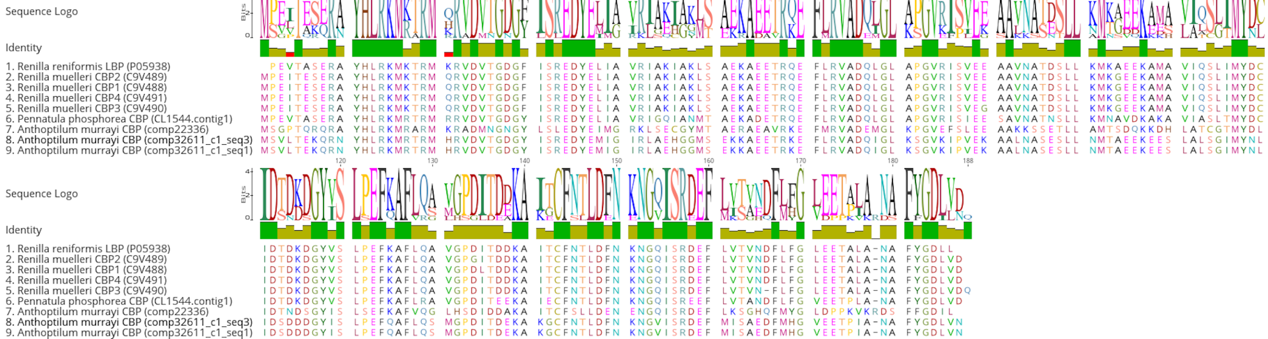

### Figure S4

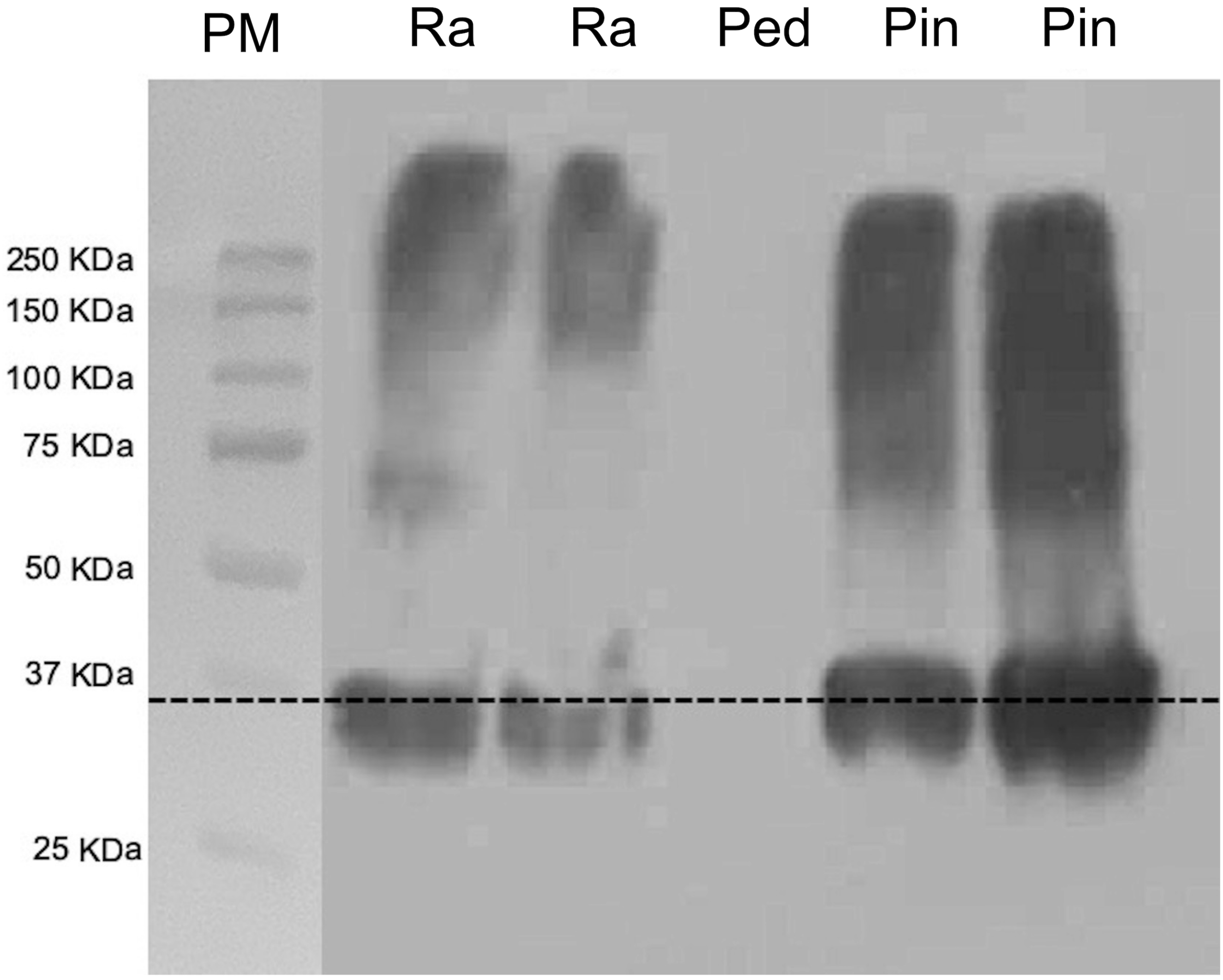

### Figure S5

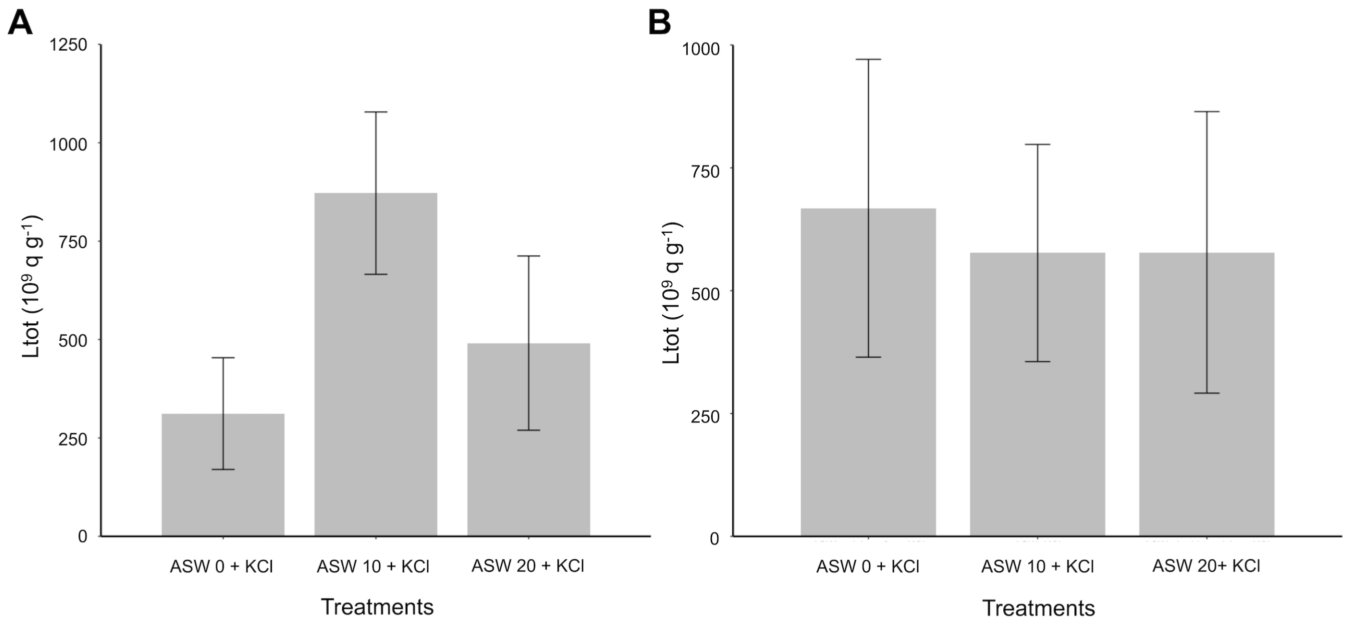
